# Complete Suspension Differentiation of Human Pluripotent Stem Cells into Pancreatic Islets Using Vertical Wheel^®^ Bioreactors

**DOI:** 10.1101/2023.08.09.552676

**Authors:** Nidheesh Dadheech, Mario Bermúdez de León, Nerea Cuesta-Gomez, Ila Tewari Jasra, Rena Pawlick, Braulio Marfil-Garza, Kevin Verhoeff, Sandhya Sapkota, Haide Razavy, Perveen Anwar, James Lyon, Patrick MacDonald, Doug O’ Gorman, Glen Jickling, AM James Shapiro

## Abstract

Advanced protocols to produce human pluripotent stem cell (SC)-derived islets show promise in functional, metabolic, and transcriptional maturation of cell therapy product to treat diabetes. Available protocols are either developed as complete planar (2D) or, in later stages, combined with suspension cultures (3D). Despite marked progress, both approaches have clear limitations for scalability, cell loss and batch to batch heterogeneity during differentiation. Using a Vertical Wheel^®^ bioreactor system, we present a highly efficient and scalable complete suspension protocol across all stages for directed differentiation of human pluripotent stem cells into functional pancreatic islets. Here, we generate homogeneous, metabolically functional, and transcriptionally enriched SC-islets and compared against adult donor islets. Generated SC-islets showed enriched endocrine cell composition (∼63% CPEP^+^NKX6.1^+^ISL1^+^) and displayed functional maturity for glucose stimulated insulin secretion (∼5-fold) during *in vitro* and post transplantation. Comprehensive stage-specific single-cell mass flow cytometry characterization with dimensional reduction analysis at stage-4 and -6 confirmed optimal maturation was achieved without heterogeneity. Notably, by 16-weeks transplantation follow-up, normal glycemic homeostasis was restored, and glucose responsive human c-peptide secretion response (2-fold) was achieved. Four months post engraftment, graft-harvested single cells displayed islet hormonal cell composition with flow cytometry, improved functional maturity by *in vivo* glucose-stimulated insulin secretion (GSIS) and enhanced transcriptional landscape with real-time expression that closely resembled patterns comparable to adult human islets. Our comprehensive evaluation of a complete suspension method applied across all stages using Vertical Wheel^®^ bioreactors for SC-islets generation highlight progressive molecular and functional maturation of islets while reducing potential cell loss and cellular heterogeneity. Such a system could potentially be scaled to deliver clinical grade SC-islet products in a closed good manufacturing practice type environment.

**One Sentence Summary:** This study describes all-stages complete suspension protocol for SC-islets generation.

## INTRODUCTION

Stem cell (SC)-derived islets offer huge potential to be a revolutionary treatment for diabetes ^1–3^. SC-islets are created by differentiating pluripotent stem cells (PSC) into insulin-producing cells, which can then be transplanted into a patient, restoring endogenous insulin production ^4, 5^. Advance methods for the optimal generation of SC-islets have generated considerable clinical interest and excitement for diabetes cell therapy ^6–8^. In fact, early success in clinical trials with embryonic stem cell (ESC)-derived islets in patients with type-1 diabetic (T1D) has reversed diabetes in small numbers of trial patients, with restoration of endogenous insulin secretion and improved glucose control ^9, 10^. SC-islet generation from ESCs solves the donor islet supply limitation for widespread islet transplantation but still relies on continuous chronic immunosuppression that raises safety concerns ^11^. ESCs and ESC-derived cell therapies have also raised ethical concerns, whereby the discovery of human induced pluripotent stem cells (iPSCs) and the opportunity to rewind adult somatic cells to regain an embryonic-like pluripotency state allows for a similar therapy option without destruction of embryos ^12^. More importantly, the generation of SC-islets from patient-derived iPSCs may one day allow for autologous transplantation without immunosuppression, thus circumventing requirement for heavy immunosuppression or immune shielding strategies to protect the graft from the recipient’s immune system ^5^.

Several investigations have presented directed differentiation of human pluripotent stem cells (PSCs) into endoderm, pancreatic progenitors and islets using matrix coated adherent plates (planar-2D) ^6, 8, 13, 14^. Alternative approaches, including one approach by our group, carried out initial planar differentiation until pancreatic progenitor cells (Stage 4, S4) followed by aggregation and transfer of re-aggregated clusters into three-dimensional (3D) suspension culture using either ultra-low attachment plates, spinner flasks, or vertical-wheel bioreactors ^15–19^. Differentiation into insulin secreting cells was achieved through the serial addition of growth factors that controlled the differentiation of the cells under 2D and 3D culture conditions. Both methods still have potential challenges in achieving high endocrine yield, functional maturity and the generation of homogeneous and safe cell products without risk of off-target growth ^11^.

Several methods have been developed to differentiate human iPSCs into islets that closely resemble primary donor islets ^7, 12, 15, 16, 20, 21^. Recent protocol optimizations work to ensure clear routes to deliver SC-islet maturation during *in vitro* differentiation through culture conditions, small molecule growth factors and inhibitors for enhanced metabolic and transcriptional maturity ^20, 22–24^. Despite these recent advancements, two main barriers remain in order to safely transplant these SC-islets in to patients with diabetes: (a) insufficient yield of differentiated products with insurmountable manufacturing costs; and (b) nonlinear transference of these protocols for scaleup platforms such as CellFactory^TM^, CellCube^®^, HYPERFlasks^®^ or multilayered CellSTACKS^®^ chambers ^25^. Harvesting high quality cells while preserving the yield of the functional product remains a persistent challenge. For clinical application, a minimum of at least ∼7,000 to 12,000 islet equivalent count (IEQ)/ Kg body weight will be required to fully reverse the diabetic state in patients ^26^. Based on estimated number of β cells, approximately a 1,000 million SC-derived β cells would be required to achieve functional glycemic control. Thus, appropriate scaling up of the process remains critical to future clinical applicability of these technologies, especially if autologous SC-products are to be a consideration. To date, development of optimized protocols capable of generating sufficient cells for autologous transplantation in patients remains a work in progress. Other protocols have generated high quality-controlled products, however, evidence demonstrating an efficient way to manufacture large batches of SC-islets through these protocols remain a challenge ^27^. One potential solution is the adaptation of suspension culture bioreactors (spinner flasks or vertical-wheel^®^ bioreactors) in SC-islet biomanufacturing. Suspension bioreactors have been shown to promote efficient generation of large masses of iPSC material ^28^ and deployment of such strategies in SC-islet GMP manufacturing might alleviate impediments.

Herein, we describe application of vertical wheel^®^ (VW) bioreactors for suspension culture across all stages of human iPSC expansion and islet-like differentiation, eliminating a need for any 2-dimensional culture step. Building on protocols described by Egli *et al* ^8, 21^, we introduced a potent cell growth inhibitor-aphidicolin, into our suspension adapted directed differentiation to eliminate contaminating residual off target cells. Patients’ reprogrammed iPSCs were inoculated as single cells and expanded as 3D clusters within VW^®^ suspension bioreactors prior to differentiation into definitive endoderm, pancreatic progenitors, and ultimately differentiated into functional islet cells. We compared the generated SC-islets to adult donor islets, in a head-to-head comparison, to quantify for their similarities and differences in cell composition, cytoarchitectural landscapes and transcriptional maturation. We conducted detailed physiological characterization of SC-islets supported by single-cell mass flow cytometry for endocrine phenotypic profiling of differentiating endocrine cells populations. This direct comparative approach to human islets for cell maturation, population heterogeneity and functional efficacy was conducted both, in vitro SC-islet differentiation and upon *in vivo* engraftment. Our comprehensive analysis shows that the generation of SC-islets from expansion through all stages of differentiation, in VW^®^ bioreactors, has the potential to increase SC-islet cell mass for scale-up while improving β cells functionality, maturity, and efficacy compared to protocols utilizing a 2D-planar differentiation protocol and/or combination protocols. Moreover, the addition of aphidicolin helped minimize off-target growth. Despite specific transcriptomic and metabolic differences between SC-islets and primary islet cells, optimal *in vivo* islet functional efficacy measured as potential to reverse diabetes in diabetic mice was accomplished.

## RESULTS

### Directed differentiation of human induced pluripotent stem cells in vertical-wheel**^®^** suspension bioreactors result in homogeneous SC-islet cluster generation

Adopting key learnings from several methods of β cells differentiation ^6–8, 15, 21^, we designed an improved seed train of 27-day complete suspension differentiation protocol for efficient and controlled differentiation of human iPSCs into pancreatic islets. We previously generated human pluripotent stem cell lines from peripheral blood mononuclear cells (PBMC) of patients with diabetes and healthy controls^28^ **(Supplementary data 1a-c).** We reported that these quality-controlled iPSC lines showed presence of pluripotency markers, viral clearance, normal karyotyping, mycoplasma absence and stem cell transcriptomics ^28^. Herein, we showed that our iPSC expansion method within the bioreactors allowed uniform sized iPSC clusters that confirmed the presence of pluripotency markers (Oct4 and Sox2) immunohistochemically and enriched populations for Oct4, Nanog, Sox2, Tra-1-60 and Tra-1-81 markers as indicated with flow cytometry and heatmap analysis (**Supplementary data 1d-f)**. Following expansion, our improved full suspension differentiation method facilitated efficient pancreatic cell differentiation of each iPSC line (n=3) in 0.1 L and 0.5 L VW^®^ bioreactor vessels. We demonstrated an efficient differentiation protocol into functional human pancreatic β cells following a six-stage differentiation (S1 to S6) (**Fig.1a)**.

**Fig. 1.**
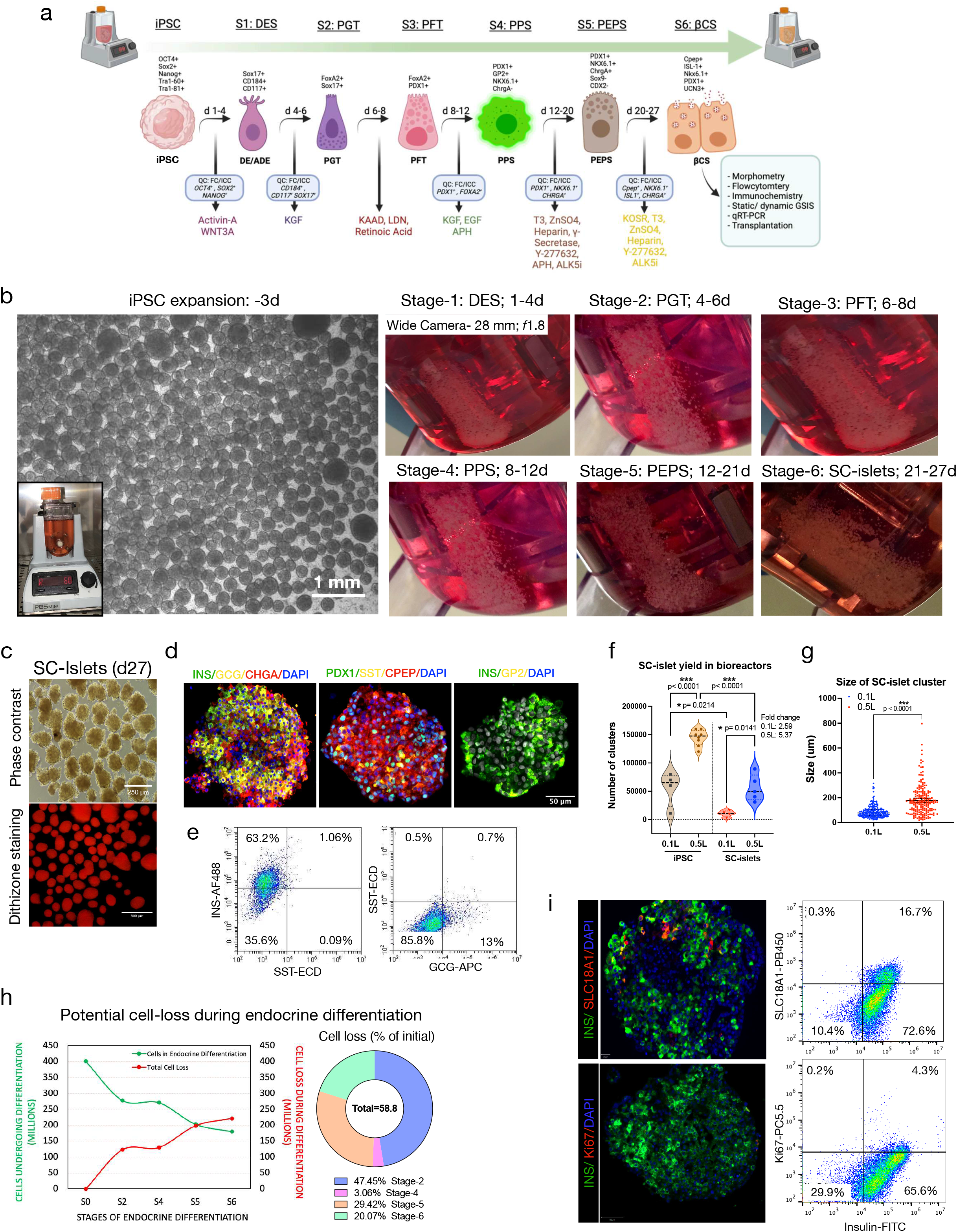
Complete suspension culture differentiation and characterization of human induced pluripotent stem cells derived islets in vertical wheel^®^ bioreactors: **a,** Overview seed-train of 27-day six-stage suspension culture differentiation protocol using VW bioreactors with quality control release parameters and stage-specific growth factors at each stage. Stages 1-7 termed as DES: definitive endoderm, PGT: primitive gut tube, PFT: posterior foregut, PPS: pancreatic progenitors, PEPS: pancreatic endocrine progenitor cell stage, βCS: beta cell stage. **b,** Phase contrast image of iPSC clusters expanded in VW bioreactor. Scale bar, 1mm. Inset shows a 0.1 L VW bioreactor with iPSCs in culture during an expansion cycle. Photographic of VW^®^ bioreactor vessel base displayed aggregated cell mass at the end of each differentiation stage between day-1 to day-27. **c,** Phase contrast and brightfield images of SC-islets for morphology and dithizone staining (red). Scale bars, 250 and 800 µm, respectively. **d,** Immunohistochemistry of SC-islets during S6 culture. Scale bars, 50 µm. **e,** Representative flow cytometry plots for islet hormonal cells-insulin, glucagon, and somatostatin. **f,** Quantification of SC-islet (S6) yield in 0.1 L and 0.5 L bioreactor vessels. One-way analysis of variance (ANOVA), multiple comparison using Turkey correction with 95% confidence interval. All data are presented as mean±sem, *n=4-8*. *\*\*\*p<0.0001*. **g,** Quantification for size of SC-islet clusters generated from 0.1 L and 0.5 L bioreactor vessels. Unpaired parametric t-test was performed with 95% confidence interval. All data are presented as mean±sem, *n=190-210*. *\*\*\*p<0.0001*. **h,** Graphical representation of cell undergoing endocrine differentiation and potential cell loss incurred during each stage from S1-S6 of islet differentiation. Parts of whole graph represents total cell loss at each stage. g, Immunohistochemistry of SC-islet clusters and representative flowcytometry for proportion of off-target SC-enterochromaffin cells (EC) and proliferating Ki-67 positive cells generation during S6.

Our cycle for iPSC expansion in a single 0.5 L VW^®^ bioreactor vessel generates 997.1 (IQR: 850-1050) million human iPSCs with uniform average clusters size of 250 µm (IQR: 125-324 µm) as previously published ^28^. Extending the expansion protocol into complete suspension differentiation for all stages of islet generation showed reproducible and homogenously sized islet mass within these bioreactor vessels at the end of each stage **(Fig.1b)**. Gravity settled cell mass at the bottom of bioreactor vessels confirmed our suspension method is efficient in preserving cellular viability and cluster cytoarchitecture, despite agitating fluid dynamics and sheer forces generated by the VW^®^ **(Fig.1b)**. Quantification for cell loss at the end of each differentiation stage revealed potential reduction in overall yield between the definitive endoderm-S2 (∼47.45%), foregut stage-S3 (∼3.06%), endocrine progenitors-S5 (∼29.42%) and terminal-S6 (∼20.07%) stages. Following S4, developing clusters maintained the yield until the end of terminal maturation S6, resulting in a final 45% cell yield. A differentiation batch starting with median 500 million iPSCs (1.7 million cells/mL; IQR: 437.5-570 cells/mL) in 0.5 L bioreactor yielded 135 million SC-islet cells (IQR: 103.3-188.2; n=5), at the end of the 27-day differentiation process **(supplementary Fig.2a)**. This protocol enabled the generation of relatively uniform and homogeneous SC-islet clusters of median size 160.6µm (IQR: 92.87-229.8µm; n=193) in 0.5 L and 77.06µm (IQR: 54.30-106.2µm; n=210) in 0.1 L bioreactor vessels. SC-islet clusters stained positive for zinc-binding dye dithizone (red) demonstrating presence of insulin storing β-like cells. Routine bioreactor culture showed median islet yield of 49,334 clusters (IQR: 35,702-78,625; n=5) in 0.5 L vessel and 10,875 (IQR: 4,688-15,263; n=4) in 0.1 L bioreactor vessel. Within these S6 clusters, we observed ∼63.21% insulin and ∼13.04% glucagon single hormonal cells, while there were ∼1.06% somatostatin cells that co-stained with insulin **(Fig.1c)**. Notably, SC-islet clusters showed presence of all islet markers-Insulin (green), Glucagon (yellow), C-peptide (red), Somatostatin (yellow), Pdx1 (green) GP2 (yellow) and Chromogranin-A (red), immunohistochemically **(Fig.1d)**. Furthermore, we observed low frequency of enterochromaffin cells marker-SLC18A1 (18.15%; SC-EC cells; IQR:16-59%-21.15%, n=4) and proliferation-Ki67 (Ki67^+^INS^+^ cells- 1.24%; IQR:0.7%-3.4%; n=4) in differentiated SC-islets that stained for insulin, out of total insulin^+^ β-like cells during S6 maturation **(Fig.1i)**.

### Cell composition analysis in bioreactor generated clusters confirms pancreatic cell generation from iPSCs

To determine the utility of VW^®^ suspension bioreactors for SC-islet differentiation, we systematically characterized cell composition from the beginning of differentiation at the definitive endoderm stage to the end of islet maturation stage using immunochemistry and flow cytometry. At definitive endoderm stage, iPSC clusters differentiated into S1 cells to show nuclear SOX17 (red) and cytoplasmic CD117 (green) co-labeled immunostaining **(Fig.2a)**. Flow cytometry analysis at S1 revealed the presence of CD184^+^CD117^+^ (94.11±1.95%; p=0.00002; n=5), and SOX17^+^CD117^+^ (83.67±5.96%; p=0.00079; n=5) cells in definitive endoderm stage **(Fig.2a supplementary data 2a)**. These cells further differentiated into CD117^+^FOXA2^+^ (79.9±9.7%; p<0.0001; n=3) cells at stage 2 and PDX1+FOXA2+ (94.2±27.1%; p=0.0035; n=3) at stage 3 prior to pancreatic progenitor differentiation **(supplementary data 2b-c)**. At stage 4, pancreatic progenitor clusters showed a uniform aggregate size of 250 µm (IQR: 146-348 µm) diameter and presence of nuclear PDX1 (red) and NKX6.1 (green) dual immunostaining **(Fig.2b)**. Flow cytometry phenotyping of S4 clusters confirmed the existence of pancreatic progenitor populations. Over 90% cells showed PDX1+ progenitor population that were also highly enriched for the pancreatic progenitor marker-GP2 (77.81±7.4%; p<0.0001; n=3), as reported previously by Nostro *et al* ^29, 30^. Of these, 67.87±7.4% cells (p=0.0001; n=3) co-expressed PDX1^+^NKX6.1^+^ with CHGA, while the remaining 23.6±7.38% population (p=0.047; n=3) were composed of CHGA+ only **(Fig.2b; supplementary data 2d)**.

**Fig. 2.**
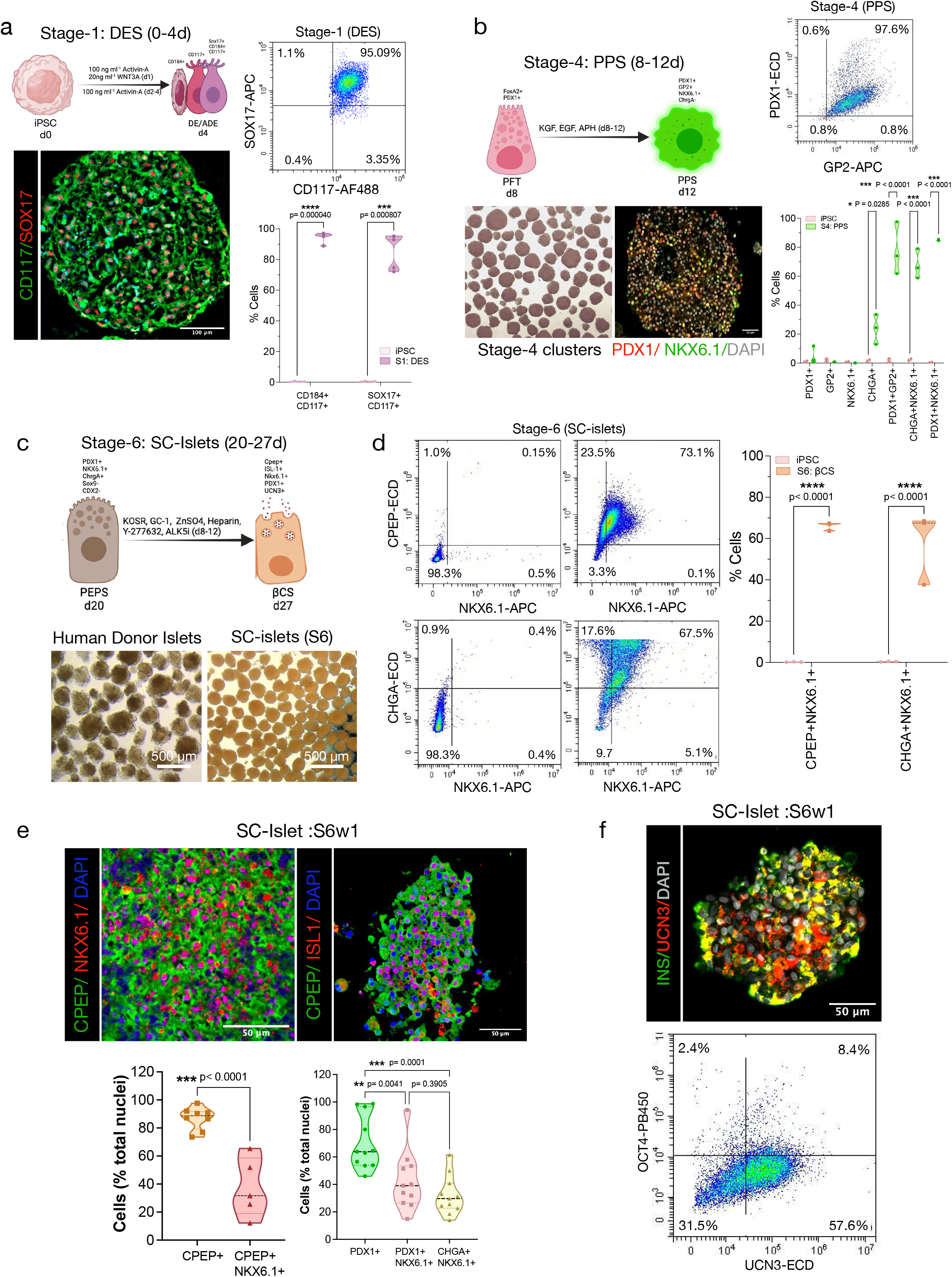
Stage-specific cell composition characterization of SC-islet cells under suspension culture differentiation: **a,** Immunohistochemistry of definitive endoderm-stage-1 cluster and representative flowcytometry for proportion of endoderm cells. Scale bars, 100 µm. Quantification by flow cytometry of CD184, CD117 and SOX17. Two-way ANOVA with multiple comparison using Šidák correction with 95% confidence interval. All data are presented as mean±sem, *n=3*. *\*\*\*\*p<0.00001* and *\*\*\*p<0.0001*. **b,** Brightfield and immunohistochemistry of stage-4 clusters and representative flowcytometry for proportion of pancreatic progenitor cells. Scale bars, 50 µm. Quantification for flowcytometry of PDX1, GP2, NKX6.1, and CHGA. Two-way ANOVA with multiple comparison using Šidák correction with 95% confidence interval. All data are presented as mean±sem, *n=3*. *\*p<0.05* and *\*\*\*p<0.0001*. **c,** Brightfield images for human donor islets and fully differentiated SC-islet clusters. Scale bars, 500 µm. **d,** Representative flow cytometry for proportion of islet cells after S6 differentiation. Quantification by flow cytometry of CPEP, NKX6.1, and CHGA. Two-way ANOVA with multiple comparison using Šidák correction with 95% confidence interval. All data are presented as mean±sem, *n=3*. *\*\*\*\*p<0.00001*. **e,** Immunohistochemistry of differentiated S6w1 clusters. Scale bars, 50 µm. Quantification for CPEP, NKX6.1, PDX1, and CHGA from immunostained clusters at S6w1. Two-way ANOVA with multiple comparison using Šidák correction with 95% confidence interval. All data are presented as mean±sem, *n=6-11*. *\*\*p<0.01 and ***p<0.0001*. **f,** Immunohistochemistry of differentiated S6w1 clusters and representative flow cytometry for proportion of maturation (UCN3) and undifferentiation (OCT4) markers. Scale bars, 50 µm.

We then characterized cells undergoing endocrine progenitor lineage differentiation and quantified endocrine committed cell populations. We identified high proportion of chromogranin-A (green) and NKX6.1(red) stained cells in S5 clusters **(supplementary data 2e),** immunohistochemically. Flow cytometry analysis for endocrine cell composition-CHGA^+^NKX6.1^+^ (64.48±3.6%; p=0.0001, n=3) cells, of which 28.1±3.5% cells (p<0.0001; n=3) co-expressed CPEP^+^NKX6.1^+^ markers **(supplementary data 2f)**. At S6, SC-islet clusters displayed islet cytoarchitecture similar to adult donor islets **(Fig.2c)**. We defined S6w0 for immature and S6w1 for mature cells in S6. Before maturation, S6w0 clusters contained 28.1% insulin expressing cells- a proportion that increased to ∼63.21% (insulin) in S6w1 clusters and ∼13.04% glucagon expressing cells. Although, somatostatin expressing remained relatively stable to ∼1.06%, these cells, however, retained insulin co-expression **(Fig.1c)**. Following maturation from S6w0 to S6w1, the proportion of cells expressing CPEP^+^NKX6.1^+^ increased to 66.10±7.2% cells (p>0.0001; n=3) of which 57.9±7.3% cells (p<0.0001; n=3) co-expressed endocrine maturation marker-CHGA. **(Fig.2e)**. Furthermore, a high proportion of SC-islet cells (∼57.64%) expressed mature hormone-UCN3. We detected ∼2.4% OCT4^+^ undifferentiated cells **(Fig.2f)**. Immunohistochemically, SC-islet cytoarchitecture at S6 stained positive for C-peptide^+^ (green) cells co-labeled with intense nuclear NKX6.1^+^ISL1^+^ (red) staining, immunohistochemically. SC-islet cell composition quantification revealed 88.7% (IQR:79.8%-71.8%; n=8; p<0.0001) cells as CPEP^+^ cells of which 31.7% (IQR:18.9%-58.6%; n=5) cells showed CPEP^+^NKX6.1^+^ staining confirming β cell-like phenotype **(Fig.2e)**. Moreover, 63.79% (IQR: 53.95-96.62%; p=0.041; n=12) of these cells were PDX1^+^, of which 39.11% (IQR: 26.49-53.41; p=0.0001; n=12) were PDX1^+^NKX6.1^+^ and 29.71% (IQR: 22.73-40.91; p=0.0001; n=12) were CHGA^+^NKX6.1^+^ **(Fig.2e)**.

### Dynamic visualization of high dimensional single-cell data detailed cellular and phenotypic heterogeneity in SC-progenitor (S4) and SC-islets (S6)

We performed high-dimensional single-cell data visualization of accutase-dispersed single cells stained with a panel of antibodies using machine learning approach **(Fig.3a**; **4a)**. Utilizing viSNE plots, we created a high-dimensional single-cell data visualisation based on the t-Distributed Stochastic Neighbor Embedding (t-SNE) algorithm from SC-derived S4 and S6 cells. viSNE helped in determining the best 2D representation of the single-cell mass flow cytometry data in terms of local and global geometry from pancreatic progenitors, differentiated SC-islet cells and islet cells. We compared cellular composition and heterogeneity between SC-derived S4 cells, S6 cells and adult islets. We observed and mapped pancreatic lineage cells into two dimensions while preserving their separation between endocrine progenitor versus SC-islet phenotypes. This allowed us to capture the heterogeneity within these populations and to highlight cell purity within the differentiation performance under optimized 3D suspension cultures against donor islets.

**Fig. 3.**
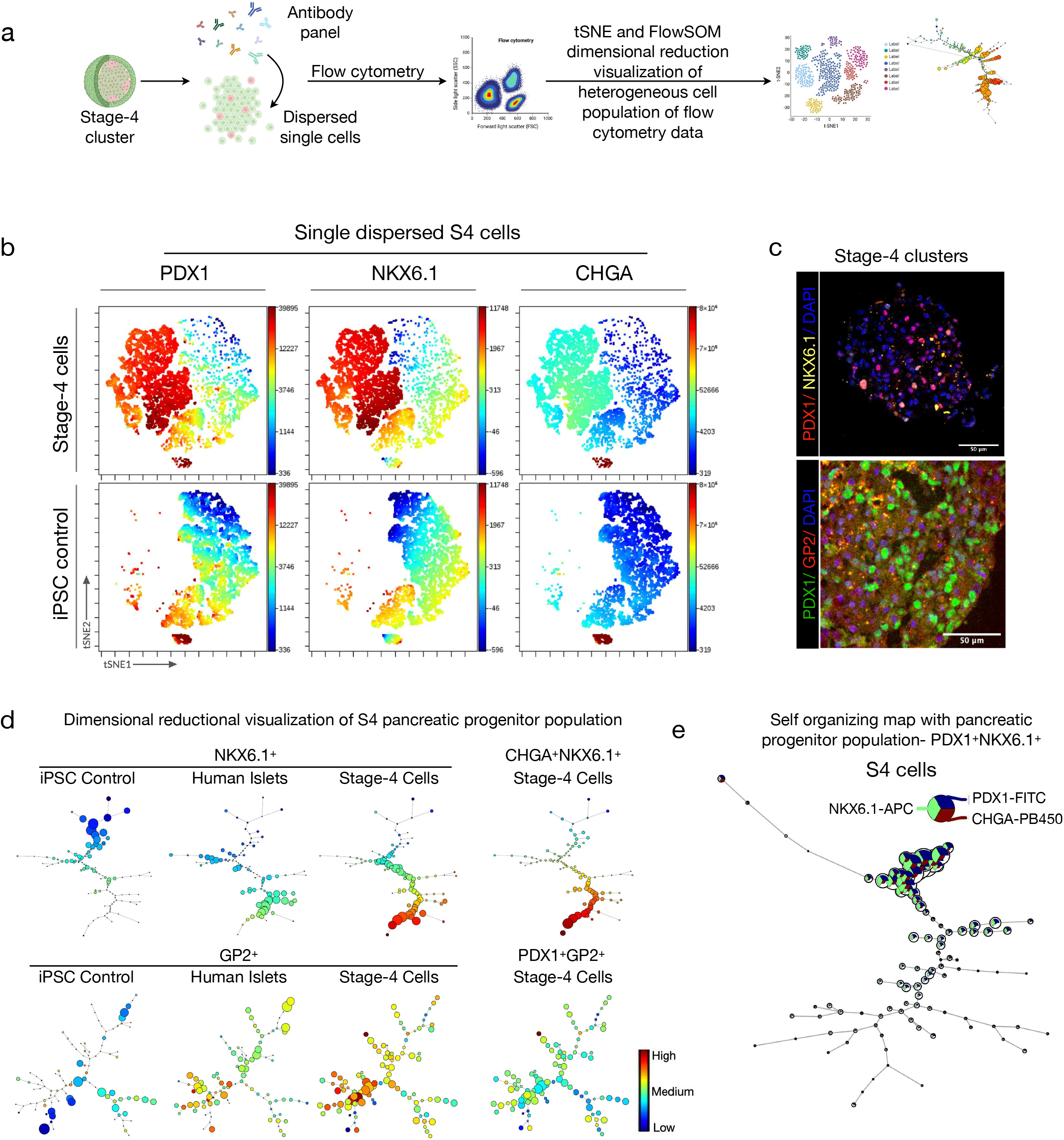
Single cell viSNE and dimensional reductional visualization for cellular heterogeneity in SC-pancreatic progenitors: **a,** Overview of experimental design to perform single cells mass flow cytometry and dimensional reductionality analysis for cellular heterogeneity in S4 cells. **b,** Single cell viSNE plot visualization of 50,000 stage-4 progenitor cells for PDX1, NKX6.1, and CHGA markers against undifferentiated iPSC control cells. Color gradient represents the density of cells from high (red) to low (blue) in viSNE islands. **c,** Immunohistochemistry of S4 clusters. Scale bars, 50 µm. **d,** Dimensional reductional visualization (FlowSOM) of the flow cytometry of S4 pancreatic progenitor cells for NKX6.1, PDX1, GP2, and CHGA against donor primary islet cells and undifferentiated iPSC control cells. Color gradient represents the density of cells from high (red) to low (blue) in endocrine lineage trajectory. **e,** Self-organizing Star Plot at single cells resolution for PDX1, NKX6.1, and CHGA in S4 cells. The minimal spanning trees of the self-organizing maps display an unsupervised clustering of the samples based on their protein expression levels (right). The heatmap scale shows the expression of each protein marker in the cell clusters (left)

**Fig. 4.**
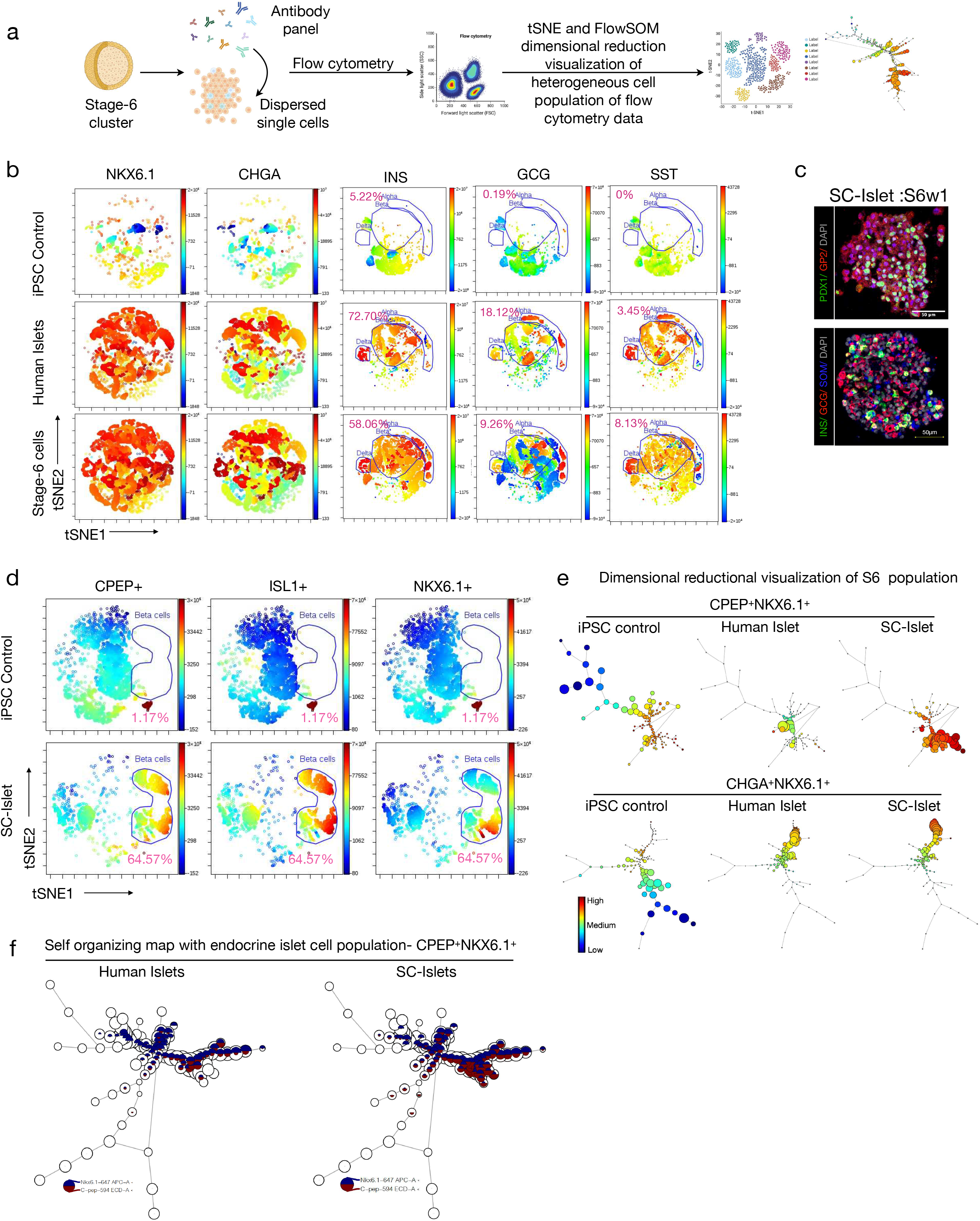
Single cell viSNE and dimensional reductional visualization for cellular heterogeneity in SC-islet cells: **a,** Overview of experimental design to perform single cells mass flowcytometry and dimensional reductionality analysis for cellular heterogeneity in SC-islets. **b,** Single cell viSNE plot visualization of 50,000 S6w1 cells for NKX6.1, CHGA and islet hormones INS, GCG, and SST against primary donor islet and undifferentiated iPSC control cells. Color gradient represents the density of cells from high (red) to low (blue) in viSNE islands. **c,** Immunohistochemistry of S6w1 clusters. Scale bars, 50 µm. **d,** viSNE representation and quantification in S6w1 islands for CPEP, NKX6.1, and ISL1aganist iPSC control. Color gradient represents the density of cells from high (red) to low (blue) in viSNE islands. **e,** Dimensional reductional visualization (FlowSOM) of the flow cytometry of S6w1 cells for CPEP and NKX6.1 against donor primary islet cells and undifferentiated iPSC control cells. Color gradient represents the density of cells from high (red) to low (blue) in endocrine lineage trajectory. **f,** Self-organizing Star Plot at single cells resolution for CPEP and NKX6.1 markers. The minimal spanning trees of the self-organizing maps display an unsupervised clustering of the samples based on their protein expression levels (right). The heatmap scale shows the expression of each protein marker in the cell clusters (left)

First, we compared progenitor marker expression on donor islets to single-cell resolution of S4 cells for phenotypic homogeneous progenitor relationship. viSNE single cell island geometry of S4 cells demonstrated comparable expression of PDX1, NKX6.1, and CHGA between adult islets, S4 and S6 cells. Mass flow cytometry viSNE analysis from 50,000 cells revealed that we were able to obtain a single-cell high resolution of cellular heterogeneity between S4 and S6 cells by multiplexing antibodies. We compared expression against representative human cadaveric islets for stochastic distribution for the pancreatic progenitor markers-PDX1, GP2, NKX6.1 and complemented with pan endocrine tracer-CHGA expression, whereas undifferentiated iPSC’s viSNE island remained negative for these markers (**Fig.3b**). Here, the intensity colour spectrum (blue-low to red-high) characterized the density of population clustered within the island cloud depicting relationship and heterogeneity. viSNE single cell island geometry of S4 cells demonstrated comparable expression of PDX1, NKX6.1, and CHGA to adult islets (**Fig.3b**). Single-cell resolution in S4 cells for PDX1-GP2 distribution revealed progenitor cell homogeneity. In addition, S4 cells metaclusters for NKX6.1 and CHGA show comparable viSNE visualization similar to donor islets **(Fig.3b)**. This was further confirmed with nuclear PDX1 (red and green), NKX6.1 (yellow), with cytoplasmic GP2 (red) immunohistochemical staining in S4 clusters (**Fig.3cf**). Collectively, our bioreactor-generated S4 product demonstrated a strongly expressing PDX1^+^ pancreatic progenitor population that also exhibited GP2^+^ and NKX6.1^+^. We then analysed the multiplexed flow data using FlowSOM tool that shows single-cell clusters that corresponds to well separated progenitor cell population and the subsets of endocrine islet populations. FlowSOM output, which concatenates the mass flow cytometry data, generated self-organizing maps to visualize clustering and dimensionality reduction. The approach has the advantage of providing a comprehensive overview of each marker’s expression level at single cell resolution and the potential to identify unsupervised cell populations. Using S4 multiplex data, the individual heatmaps that are projected onto the self-organizing maps for each marker show the expression level of each progenitor markers for all cell subpopulations **(Fig.3d)**. Evidently, NKX6.1 was uniformly high (red) in all subpopulations of S4 cells and medium in donor islets compared to iPSC control. GP2 expression was also detected at high to medium (red - yellow) in S4 cells, similar to donor islets but negative in iPSC control. Moreover, combined detected of high NKX6.1^+^CHGA^+^ and medium PDX1^+^GP2^+^ expression was uniformly identified across all cell subpopulation of S4 clusters **(Fig.3d)**. Notably, the extent of pancreatic progenitor population within the suspension differentiation of S4 clusters was pronounced S4 in self-organizing maps. FlowSOM analysis showed good overlap in signals for PDX1^+^GP2^+^, thereby providing added validation for homogeneous pancreatic progenitor population during differentiation **(Fig.3e)**.

The same approach was used to evaluate the cellular relationships and cellular heterogeneity and composition between (S6) SC-islets, iPSCs and donor islets **(Fig.4a)**. The panel of endocrine antibodies distinguished and separated hormone and endocrine populations. SC-islet cells exhibited intense and uniform NKX6.1 and CHGA expressing population distribution, similar to adult donor islets, but not found in iPSCs **(Fig.4b)**. Here, viSNE maps showed three distinct metaclusters-GCG^+^, INS^+^, and STT^+^ in S6 SC-islet cells, similar to donor islets **(Fig.4b**). A high-dimensional single-cell analysis of islet hormones expression revealed that SC-islets had undergone endocrine (islet) reprogramming. We gated endocrine subtypes in SC-islet clusters using viSNE plots to assess the hormone-expressing cell composition and compared against primary islets. To validate the map, we used an independently derived classification of the cells to islet subtypes, based on manual gating of a series of biaxial plots (see Materials and Methods). While viSNE was not provided with this classification or any knowledge of non-endocrine cells subsets, it grouped cells in the same subsets together and separated the subsets from one another based on distribution of islet-endocrine markers. viSNE accurately distinguished NKX6.1^+^ and CHGA^+^ (93%; IQR: 92.25-94.58% and 90.22%; IQR: 85.53-94.97%; n=4 respectively) cells as mature endocrine cells in S6 similar to adult donor islets, but not in undifferentiated iPSCs **(Extended data-5)**. We identified 48.6±5.0% (p=0.0026, n=3) INS^+^ β cells, 9.6±0.5% (p=0.0018, n=3) GCG^+^ α cells and 6.2±1.6% (p=0.0024, n=3) SST^+^ (δ cells) S6 differentiated SC-islets from three independent donor iPSC lines **(Fig.4b and Extended data-4)**. Notably, INS^+^ β cells formed a distinct subset in the viSNE island distinct from GCG^+^ α cells **(Fig.4b)**. In parallel, immunohistochemical staining for β cell-specific PDX1 (green), GP2 (red), and islet hormones INS (green), GCG (red) and SST (blue) endorsed an efficient endocrine programming for SC-islet generation in suspension bioreactors **(Fig.4c)**.

Further, mass flow cytometry provides a unique opportunity to interrogate large numbers of cells with multiple parameters at the single-cell level in parallel, thus facilitating identification of rare cell types and subtype-specific behavior ^31, 32^. To take advantage of this approach, we performed viSNE analysis on SC-islets to exclusively gate for β cells using surrogate antibody for gold standard β cell markers-CPEP, NKX6.1 and ISL1. This process confirmed the presence of true β cells within the SC-islets. Using spectral maps as a guide, we hand-delineated β cells cluster to further detect for co-expression with NKX6.1 and ISL1 **(Fig.4d)**. We performed a number of analyses to evaluate the robustness of viSNE. The viSNE map in Fig.3 and 4 includes 10,000 cells that were subsampled from the complete data set (each for iPSC, -S4, -S6 and donor islets). We independently subsampled multiple subsets of the data and ran viSNE on each, resulting in similar viSNE maps that conserve the separation between subsets (**Extended data-5 and 6)** This supports the notion that the viSNE map is consistent and reliable in representing an optimal islet cell composition.

To test the efficiency of islet differentiation homogeneity, FlowSOM enabled defined dimensional resolution of each islet cell subtype based on islet-specific marker expression. Population trajectory in self-organizing map from S6 cells showed marked increase in expression (red) of C-PEP and NKX6.1 markers, compared to human donor islets. Similarly, the meta clusters of all cell subpopulations from S6 and donor islets exhibited comparable high-medium expression of CHGA and NKX6.1 **(Fig.4e)**. Collectively, FlowSOM heatmaps from S4 and S6 cells described the distribution of cell populations and their expression levels for pancreatic progenitor, pancreatic endocrine progenitor, and hormone secreting markers. Remarkably, minimal spreading of subpopulation in the self-organizing maps reflected limited cellular heterogeneity in cell compositions.

### Suspension SC-islets transcriptionally mature *in vitro*

To investigate the transcriptional maturation and changes associated with *in vitro* iPSC to SC-islet differentiation in suspension culture bioreactors, we performed endocrine transcriptomic analysis using Taqman Low Density Genes Array (TLDA) cards. We compared the transcriptome of SC-islets to datasets from adult islets and undifferentiated iPSCs for forty-eight critical pancreatic differentiation pathway-specific genes encoded in low density array cards. Gene array configuration can be found in **table S7**. The endocrine gene datasets were also assessed at every stage of islet differentiation. Despite minor differences after normalization with iPSC control, heatmap transcriptome clustering analysis for forty-eight genes from three independent SC-islet (S6w1) batches was comparable to three primary human donor islets **(Fig.5a)**. Real-time transcriptomic data show a log2 fold change gene expression matrix from cells using a colour gradient module (-1.5: blue to 1.5: red) and clustered with the complete clustering method. All three batches of SC-islets, donor islets, and undifferentiated iPSCs were efficiently clustered together using a heatmap matrix, where SC-islets and donor islets had a close interrelationship. The transition of relative gene expression of endocrine genes from S1 to S6 revealed a sharp upregulation of *INS* transcripts beginning in S4 and continuing until S6. The maturation marker *UCN3* and the endocrine transcript *ISL1* increased between S4 and S6 **(Fig.5b)**. Furthermore, the enterochromaffin lineage marker *SLC18A1* also increased throughout iPSC differentiation **(Fig.5b)**. Notably, with the exception of *SLC18A1*, the level of each endocrine transcript in S6w1 clusters was found to be equivalent to that of donor islets. The statistical comparison for each gene amongst multiple groups are described in **Extended source data table 1**.

**Fig. 5.**
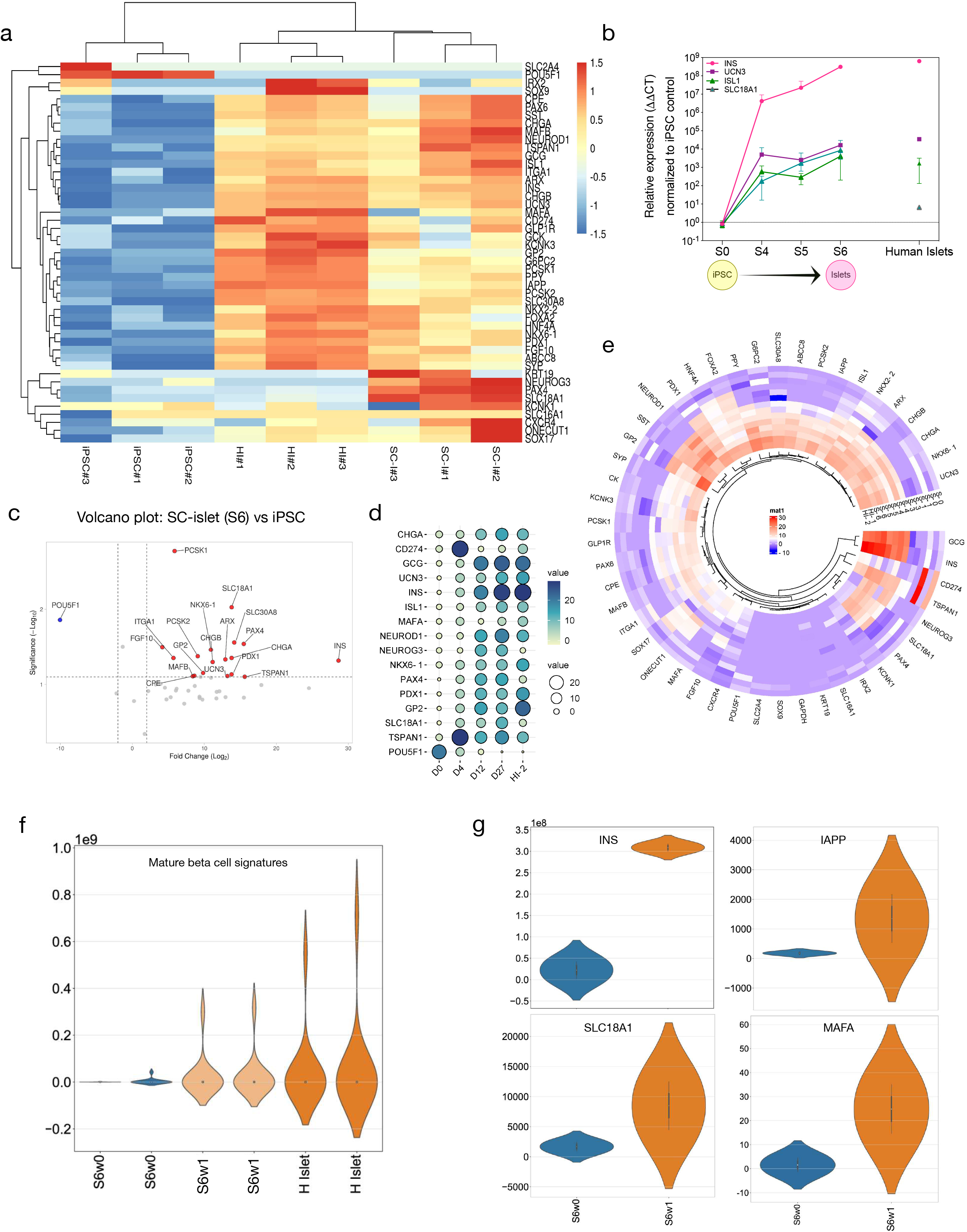
Transcriptional maturation of stem cell-derived islets: **a,** Heat map projection of gene clusters for islet differentiation in SC-islets compared to primary islets and undifferentiated iPSCs. **b,** Real time expression of selected genes upregulated in cells undergoing differentiation from iPSCs to S6. **c,** Volcano plot visualisation of SC-islets against undifferentiated iPSCs to show key gene sets upregulated and downregulated in vitro. **d,** Bubble plot projection (left) of key islet-specific gene sets upregulated over time during differentiation stages progression. Heatmap color scale (top right) show level of gene expression and buddle size indicator (bottom right) represents density of each transcript during stages of differentiation. **e,** Circular cluster heatmap projection shows average expression of all genes associated with pancreatic differentiation in time course from stages 0, 1, 3, 4, 5 and 6 against primary donor islets, *n=2* each stage. Heatmap scale show level of gene expression (>0 upregulation and <0 downregulation). **f,** Violin plots representing mature beta cell signature in SC-islets from S6w1 cells compared to adult human islets and immature S6w0 cells. **g,** Violin plots representing level of expression of genes associated with mature beta cell hallmark processes in S6w1 cells from time of origin.

A volcano plot visualisation of significantly upregulated genes transcript between S6w0 (immature) and S6w1 (mature) cells revealed increased expression of many maturation-related genes including *MAFB, PCSK1/2, G6PC2*, *ITGA1, SLC30A8 (ZnT8), CPE*, and *TSPAN1*. Furthermore, comparison of S6w1 transcriptome to iPSC transcriptome showed the upregulation of multiple genes associated with β cell identity and insulin secretion including *INS, PDX1, UCN3, NKX6.1, GP2, CHGB, NGN3, NEUROD1, MAFB*, and *PAX4*; as well as upregulation of α cells associated genes like *ARX, CHGB, ISL1 and MAFB*. Furthermore, the pluripotency associated gene *POU5F1* was downregulated in S6w1 cells compared to S6w0 **(Fig.5c)**.

We then focused on the SC-islet transcriptome during the differentiation trajectory to determine the transcriptional changes associated with differentiation and functional maturity. Bubble plot correlation analysis indicated that S6w1 cells (d27) and donor islet cells were transcriptionally more similar among each other than S5w0 (d12) cells. We compared the average gene expression of known cell maturation markers at d0, 4, 12, and 27. Expression of *INS, GCG, UCN3, NEUROD1, MAFA,* and *CHGA* show increase at d4 until d27, whereas other progenitor markers such as *NEUROG3, PDX1, PAX4,* and *GP2* were expressed throughout differentiation. Interestingly, *CD274 (PDL1)* and *TSPAN1* were initially upregulated but their expression decreased at later stages to levels comparable to mature donor islets. *POU5F1* levels at day 27 were comparable to donor islet cells **(Fig.5d)**.

We investigated the temporal gene regulation of endocrine genes throughout the entire islet developmental pathway, from iPSC (S0) to SC-islets (S6). A heatmap comparison of the islet transcriptome throughout S6 cell differentiation showed transcriptional upregulation of pancreas development associated genes **(Fig.5e)**. Transcript levels were graded using a colour intensity gradient, with significantly upregulated genes scored in red and differentially repressed genes scored in blue. Endocrine patterning gene clustering suggested progressive transcripts upregulation of *INS, GCG, SST* and *PPY*. S6 iPSC-islets had upregulated genes associated with pancreatic differentiation such as *NKX6.1, CHGA, NKX2.2, ISL1, FOXA2, PDX1, NEUROD1*, and *GP2* throughout stages 3 to 6. Furthermore, maturation-related genes such as *UCN3, CHGB, IAPP, PCSK8, ABCC8, SLC30A8, G6PC2, KNCK3, PCSK1/2, GLP1R, CPE, MAFB*, and *ITGA1* were upregulated at S5 and S6. Non-endocrine and side population transcript levels such as *ONECUT1,* and *POU5F1* were effectively repressed during terminal maturation **(Fig.5e)** while *SOX9* and *KRT19* remained lower and unchanged, indicating that the extent of transcriptional maturation was uniformly endocrine.

Next, we investigated the transcriptional changes specifically associated to β cell signatures. We compared immature S6w0 cells (prior to maturation stage), fully developed S6w1 cells against primary donor β cells. Violin plot visualization of β cell specific maturation gene matrix including *INS, UCN3, MAFA, CHGB, IAPP, PCSK8, ABCC8, SLC30A8, G6PC2, KNCK3, PCSK1/2, GLP1R, CPE, MAFB*, and *ITGA1* showed that S6w1 cells had equivalent expression to primary donor cells **(Fig.5f)**. Individual violin plots between S6w0 and S6w1 cells show that *INS, IAPP, SLC30A8*, and *MAFA* transcripts are significantly upregulated **(Fig.5g)**.

### Suspension SC-islets show functional insulin secretion machinery

Human islets exhibited tightly regulated biphasic insulin secretion in response to changes in glucose concentration. We performed static and dynamic glucose responsive insulin release assays to understand the functional capacity of SC-islets and the efficiency of their functional machinery for glucose sensitivity. While the static profiles generated marked insulin release in response to glucose, the dynamic perifusion of SC-islets showed immature insulin secretion in response to glucose, the GLP-1 analogue Exendin-4 (25-35 minutes) and KCl depolarization **(Fig.6a and Extended data-1)**. Static incubation of S6w1 islets displayed a significant increase in insulin secretion with fold change of 4.67±2.8 (p=0.0269; n=5) after exposure to high glucose (16.7 mM) and 6.71±4.8 (p>0.0001; n=5) after membrane depolarization with KCl (35 mM) **(Fig.6b)**.

**Fig. 6.**
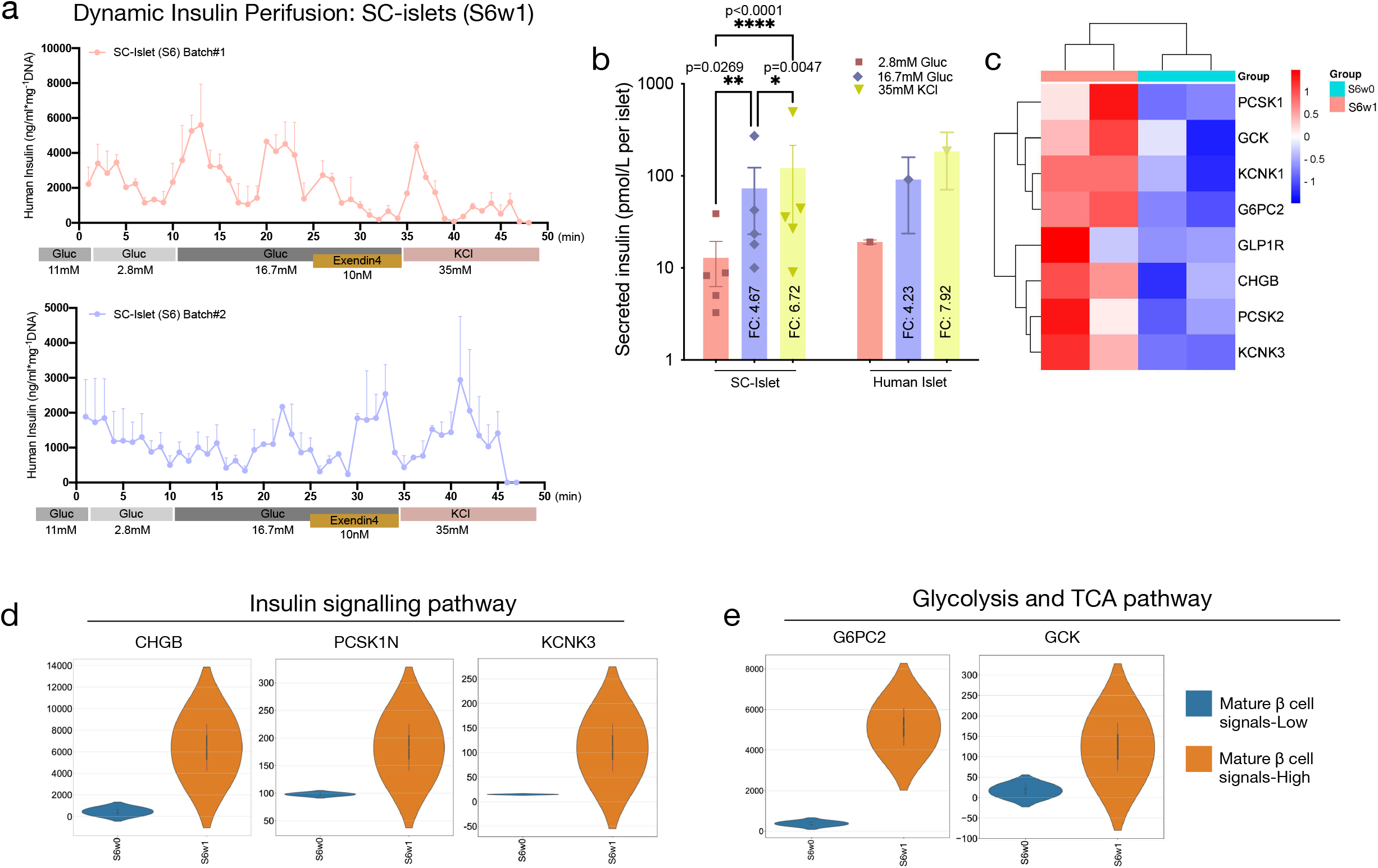
Functional phenotyping of stem cell-derived islets: **a,** Insulin secretion response to dynamic perifusion with 11 mM (G11), 2.8 mM (G3), 16.7 mM (G17) glucose, 10 ng/ml exendin-4 (EX4), and 35 mM KCl. Secretion tracing was normalized to total DNA content *(n=3)* from fifty S6w1 clusters from two independent differentiation batch replicates. **b,** Insulin secretion response to varying glucose concentrations from G3 to G17 and KCl in static incubations from S6w1 clusters in comparison to adult donor islets. Normalized to per islet tested for release assay, *n=5*. Average secretion of technical replicates, *n=2*, plotted for adult donor islets. Two-way ANOVA with multiple comparison using Šidák correction with 95% confidence interval. All data are presented as mean±sem, *n=5*. *\*p<0.05, **p<0.001*, and *\*\*\*\*p<0.00001*. c, Heatmap projection of gene expression profile in S6w1 cells against immature S6w0 cells for genes associated with mature beta cell hallmark processes, glucose metabolism and functionality. **d-e,** Violin plots representing average expression of genes responsible for insulin secretion, oxidative phosphorylation, glycolysis, and TCA cycle pathways.

We then investigated the transcriptional changes that enabled SC-islet in vitro functional maturity. Several regulatory genes associated with β cell physiological and functional maturation were found upregulated in terminally differentiated S6w1 cells. Our data indicated that glucose control regulating genes (*PSCK1/2, GCK, IAPP, G6PC2*) along with exocytosis controlling genes (*KCNK1, GLP1R, CHGB*) were all upregulated in S6w1 cells compared to immature S6w0 cells. Heatmap cluster matrix analysis of maturation-associated upregulated (red) genes in SC-islets from two independent batches of S6w1 islets clustered together than immature S6w0 expression level (blue) **(Fig.6c)**. Transcription factors involved in insulin signalling (*PCSK1N, CHGB, KCNK3*) pathway were promoted. Glycolysis and TCA pathway related genes-*G6PC2, GCK* were upregulated during S6w1 β cells maturation, along with genes involved in transportation process that facilitate insulin exocytosis (*SLC2A4, SLC30A8*) **(Fig.6d-e)**. By contrast, expression of disallowed gene monocarboxylate transporter *SLC16A1 (MCT1)* was reduced during *in vitro* maturation of S6w1 cells (**Extended data-7)**. These findings are consistent with our functional and molecular characterization. These transcriptional changes suggest that suspension generated S6w1 cells have improved insulin production and active exocytosis machinery, which are all hallmarks of maturing β cells as they develop.

### Human pancreatic progenitors and SC-islets normalize hyperglycemia in mice after implantation

To investigate the *in vivo* functional efficacy of immature pancreatic progenitor cells (S4) and mature islet cells at S6w1, we transplanted the cell products under the kidney capsule of streptozotocin (STZ) induced diabetic immunodeficient SCID beige mice **(Fig.7a)**. Prior to transplantation, cell clusters from each batch were characterized with flow cytometry. Over 84.89±0.3% cells of S4w0 were PDX1^+^NKX6.1^+^ while 66.10±1.3% cells of S6w1 were CPEP^+^NKX6.1^+^ **(Fig.7b)**. Histological assessment from harvested grafts after 3-months post transplantation of S4 and S6 cells, showed endocrine tissue morphology underneath the kidney capsule. Immunochemical staining for human insulin on sliced and paraffin fixed grafts confirmed the presence of insulin positive (red) SC-β cells **(Fig.7c)**. We monitored glucose levels of the transplanted mice over 180 days to access graft engraftment, *in vivo* differentiation, and functional efficacy with long-term graft maturation. Non-transplanted STZ-induced diabetic control mice (red) exhibited elevated glucose levels >20 mM over 90 days **(Fig.7d)**. Mice transplanted with S4 clusters (blue) displayed gradual reversal of hyperglycemia over 60 days post implantation and continue to reverse until 150 days. Interestingly, S6w1 transplanted mice (green) displayed rapid decline in elevated blood glucose within 42t (S4: p=0.0015, n=13 and S6: p=0.00020, n=19) days post implantation that remain persisted until 105 days where S6 cells exerted stable control of blood sugar significantly higher than S4 cells (p=0.0087) **(Fig.7d)**. Notably, area under the curve for blood glucose measurements indicated faster rapid glycemic reversal with fully differentiated SC-islets **(Fig.7e)**. Transplanted mice with S4 cells showed detectable human c-peptide secretion response at 2 months post-transplantation; c-peptide secretion increased over 2-folds at 3 months post-transplantation **(Fig.7f)**. Likewise, S6w1 transplanted mice exhibited stimulated human c-peptide secretion in the blood stream within 2 months, which increased by 2-fold at 3- and 4-months post-transplantation **(Fig.7g)**. Additionally, intraperitoneal glucose tolerance test in these mice revealed improved glucose control at 3- and 4-months post-transplantation compared to 2 months post-transplantation in S4 and S6 implanted animals, respectively. Area under the curve for S4 implanted cells show modest effect in glucose tolerance than naïve mice whereas S6 mature cells in mice at 4 months implantation displayed significantly improved glucose tolerance than S4 cells and naïve animals at 90 and 120 min (p=0.0062 and 0.020 respectively) time points **(Fig.7h-i)**.

**Fig. 7.**
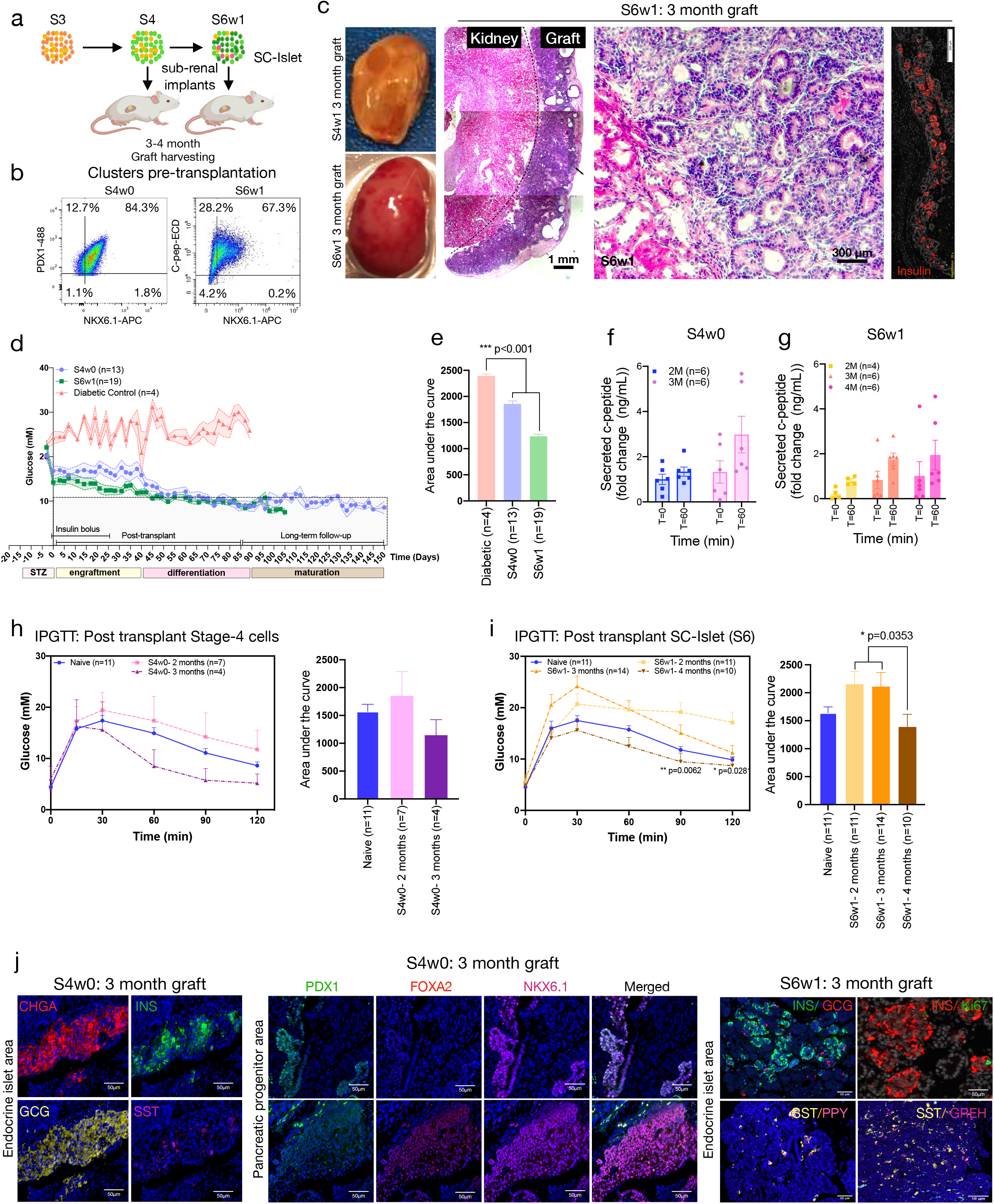
*In vivo* transplantation of stem cell-derived pancreatic progenitors and islet clusters: **a,** Experimental overview, and timeline representation for graft assessment in S4 and S6 cells, post 3-4 months transplantation in mice. **b,** representative flow cytometry confirmation for SC-S4/S6 cell composition, prior to transplantation. **c,** Photographs of harvested kidneys from S4w0 and S6w1 transplanted mice after 3 months of engraftment. Representative images display gross histology (hematoxylin and Eosin) and immunohistochemistry of endocrine graft region under renal subcapsular space from 3 months transplanted mice tissues. Scale bars, 1 mm and 200-300 µm. **d,** Quantification for *in vivo* efficacy of S4w0 vs S6w1 cells to correct elevated blood glucose in mice until 150 days post transplantation. Two-way ANOVA with average mixed effect using Turkey correction with 95% confidence interval, at multiple time points comparison. All data are presented as mean±sem, *n=13-19* mice, *ns= p>0.05.* **e,** Quantification for area under the curve to show efficient glucose lowering effect from S4w0 and S6w1 cells for 1-3 months post transplantation. One-way ANOVA and multiple comparison using Turkey correction with 95% confidence interval. All data are presented as mean±sem, *n=4-19*, *\*\*\*p<0.001.* **f-g,** In-vivo glucose stimulated human c-peptide secretion from S4w0 and S6w1 transplanted mice at 2-, 3- and 4-months post transplantation. All data are presented as mean±sem, *n=6.* **h-i,** Quantification of *in vivo* functional efficacy of impaired glucose tolerance at 2-, 3-, and 4-months post transplantation in S4w0 and S6w1 engrafted mice. Bar graphs represent area under the curve to show potential effect in glucose tolerance and graft function. Two-way ANOVA and multiple comparison using Šidák correction with 95% confidence interval was performed. All data are presented as mean±sem, *n=4-11* mice, *\*p<0.05* and *\*\*p<0.001*. j, Immunohistochemistry of harvested graft tissues from S4w0 and S6w1 implanted mice after maturation at 3 months post transplantation. Scale bars, 50 µm.

Retrieved tissues at 3 months post implantation were assessed for islet hormones, endocrine fraction, and progenitor cell composition using immunohistochemistry. Grafts retrieved at 3 months post-transplantation from the mice transplanted with S4 cells showed presence of islet hormones including β-like cells (insulin-green), α-like cell (glucagon-yellow), δ-like cells (somatostatin-magenta), and chromogranin-A cells (red). Few β-like cells co-labeled with α-like cell suggesting immaturity or polyhormonality **(Fig.7j)**. A marked proportion of S4 grafted cells retained pancreatic progenitor identity characterized as co-staining for PDX1 (green), FOXA2 (red) and NKX6.1 (magenta) in multiple regions of the harvested graft tissues. Conversely, S6w1 implanted cells showed endocrine cell maturation of the grafted tissues in all islet hormone cells. S6w1 graft sections showed distinct immunostaining for β-like cells (insulin-green), α-like cell (glucagon-red), δ-like cells (somatostatin-yellow), γ-like cells (pancreatic polypeptide-magenta) and grehlin cells (magenta). At 3 months, marginal β-like cells were co-labeled with Ki67 (green) and insulin (red) immunostaining, suggesting proliferative potential in SC-β cells post transplantation **(Fig.7j)**. Collectively, S6w1 cells showed improved in-vivo differentiation potential and enhanced functional maturation for rapid diabetes reversal than stage 4 cells.

### Graft-derived SC-islet single cells show improved endocrine composition with enhanced transcriptional and functional maturity

To assess the endocrine cell composition within the harvested grafts and investigate the transcriptional changes associated with *in vivo* maturation, we performed single cell dispersion of graft-harvested tissues at day 120. Single cells were characterized with flow cytometry and with TaqMan gene arrays. Furthermore, we re-clustered the single cells, and we performed glucose stimulated human insulin secretion assay in the re-aggregated graft cells **(Fig.8a)**. We obtained a dataset comprising from 10,000-50,000 single graft cells and clustered according to origin and cell identity based on flow cytometry. The entire graft cell composition from transplanted animals (n=5) revealed that ∼77% of the cells were of endocrine origin while the ∼23% of remaining cells were from other non-pancreatic origin. Within the endocrine population, ∼44% and ∼24% of the cells had endocrine and pancreatic progenitor phenotype, respectively. Furthermore, ∼32% of the cells depicted β-like cell phenotype of which nearly 6% were proliferating β-like cells **(Fig.8b)**. This *in vivo* endocrine maturation trajectory was highly evident from a comparative profiling of graft harvested and S6 cells for true SC-β cells versus SC-EC cells commitment pre- and post-implantation. Notably, 31.3±8.0% (p=0.0070; n=4) of graft matured SC-β cells at d120 acquired β cell-like phenotype, in vivo, while losing SLC18A1 expression only 5.04±1.3% (p>0.0001, n=4) cells retained INS^+^SLC18A1^+^ expression. S6 cells, pre-transplantation, inherited 18.6±1.2% INS^+^SLC18A1^+^ with majority of 58.4±2.9% (p=0.002, n=4) cells as INS+ expressing **(Fig.8c)**.

**Fig. 8.**
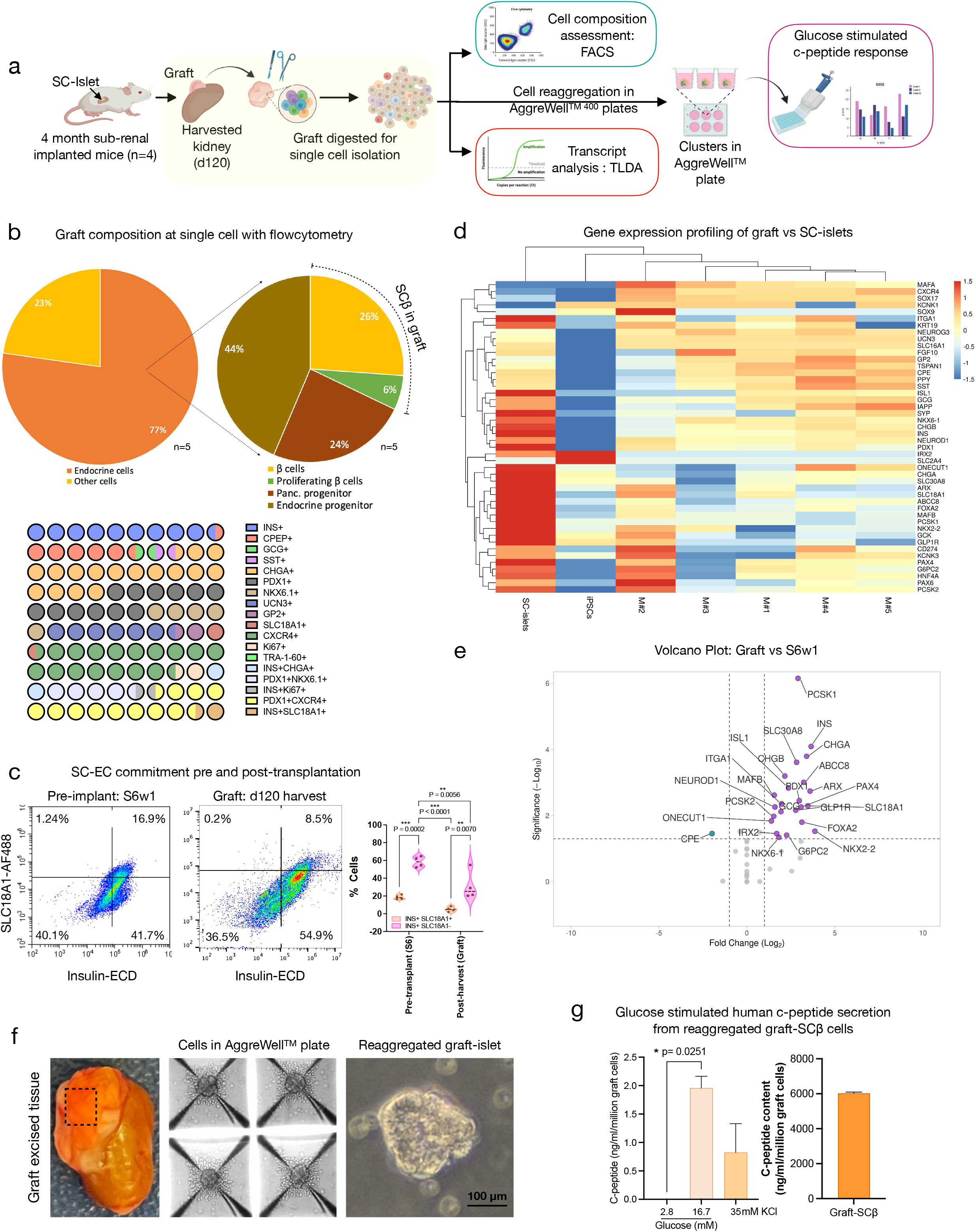
Graft-harvested cell characterization post transplantation and maturation: **a,** Experimental overview for cell composition, transcriptomic and functional characterization from graft matured and harvested SC-islet cells. **b,** Flow cytometry-based graft cell composition pi-chart quantification and representation for classification of multiple pancreatic cell populations. Heat map (below) shows parts of whole representation of entire graft cell composition. *n=4-7* mice. **c,** Representative flow cytometry plots for off-target SC-EC cell characterization and quantification pre and post transplantation. Two-way ANOVA and multiple comparison using Šidák correction with 95% confidence interval was performed. All data are presented as mean±sem, *n=4*, \**\*p<0.01* and *\*\*\*p<0.0001*. **d**, Heat map projection of gene clusters for islet differentiation and maturation in SC-islets implanted graft-harvested tissues in comparison to S6w1 islets (average of 3 independent differentiation preps) pre-transplantation and undifferentiated iPSCs (average of 3 iPSC lines). Color scale (top right) represents gene upregulation (red; >0) and downregulation (blue; <0), Graft assessment form *n=5* mice. SC-islet or iPSC datasets are represented as an average of *n=3* independent replicates. **e,** Volcano plot representation shows extent of endocrine maturation 3-months post engraftment with critical beta, alpha, and endocrine maturation associated genes upregulation against S6w1 cells, pre-transplantation. **f,** Image of excised S6w1 graft from harvested kidney, 3 months post transplantation. Reaggregation and pseudo islet formation of graft harvested single dispersed cells in AggreWell400^TM^ plate. Representative image of a reaggregated graft-derived SC-islet cluster. Scale bars, 100 µm. **g,** Quantification for *in-vitro* glucose stimulated human c-peptide secretion (left) and total human C-peptide content from one million graft harvested cells reaggregated into islet-like clusters. Two-way ANOVA and multiple comparison using Šidák correction with 95% confidence interval was performed. All data are presented as mean±sd, *n=3 technical replicates*, \**p<0.05*.

We then looked for transcriptomic maturation of graft harvested S6w1 cells after 120 days of maturation. Gene array quantification for 48 islet maturation genes revealed significantly efficient endocrine maturation for islet hormonal cells (descriptive statistics for each gene is shown in **Table-2**). Comparative analysis of datasets from 100,000 single dispersed graft-harvested cells from transplanted mice displayed enriched islet transcriptome and upregulation of key differentiation genes. Heatmap clustering matrix confirmed upregulated of *MAFA, KCNK1, NGN3, UCN3, TSPAN, IAPP* and *CHGB*) compared to differentiated S6w1 cells prior to the transplant. Heatmap confirmed downregulation of off-target genes: *SOX9, KRT19, POU5F1* **(Fig.8d)**. A volcano plot comparison of datasets from graft-harvested cells against S6w1 (before implantation) confirmed the upregulation of multiple islet population genes for (β, and α) and functional maturation associated genes. Genes for β cells: *INS, PDX1, CHGB, ISL1, GLP1R, NEUROD1, MAFB,* and *PAX4*; α cells*: ARX, CHGB, ISL1, MAFB*; endocrine maturation genes: *MAFB, PCSK1, ITGA1, SLC30A8 (ZnT8), ABCC8 (SUR1)* were significantly upregulated. **(Fig.8e)**.

To test the extent of functional maturity and insulin secretion potential of graft-harvested SC-islets, we challenged the re-aggregated pseudo clusters (in Aggrewell400^TM^ plates) for glucose stimulated human c-peptide secretion **(Fig.8f)**. A 1.96 ng/ml (IQR:2.1-1.8 ng/ml; n=2 technical replicates) of human c-peptide secretion in response to high glucose (16.7 mM) exposure was measured from 30 clusters comprised of one million graft-retrieved cells. The total c-peptide content from pseudo islets was 6,000 ng/ml (IQR:6028) **(Fig.8g)**. Collectively, our results indicated that suspension (bioreactor) differentiated SC-islets (S6w1) optimally engrafted and underwent transcriptional and functional maturity post transplantation to reverse diabetes in mice.

## DISCUSSION

The differentiation of human pluripotent stem cells into pancreatic islets is a promising approach for developing cell-based therapies for diabetes. However, achieving functional maturation of these cells *in vitro* remains a challenge. One method that has shown promise for improving the differentiation efficiency and functionality of stem cell-derived islets is the use of bioreactors ^15, 20, 24^. VW® bioreactors are a type of bioreactors that provide a highly controlled and dynamic environment for cell growth and directed differentiation with minimal sheer stress^28, 33, 34^.

In this study, we describe a complete suspension protocol to expand human induced pluripotent stem cells and further generate SC-islets using VW® bioreactors with high efficacy, functionality, and reproducibility. The generated clusters are homogeneous and exhibited defined endocrine cell composition, glucose regulated insulin release, and transcriptional maturation similar to that of primary human donor cadaveric islets. The utility of VW®-based bioreactor system enhanced the differentiation efficiency and functionality of the SC-islets compared to standard planar culture conditions as represented by cell composition through flow cytometry, immunostaining for several islet markers and *in vivo* functional parameters. In particular, the bioreactor system promoted the formation of large numbers and more uniform islet-like clusters, which displayed glucose-stimulated insulin secretion and expression of key β cell markers at the end of stage 6. Comparatively, the suspension (3D) generated S6 clusters showed dynamic glucose control and efficient insulin secretion machinery similar to primary islets. These findings are in accordance with several previously published and optimized protocols for SC-islet generation in 2D planar^6–8, 13, 14^ or combined with suspension cultures^15, 35^.

Despite the potential cell loss (∼58%) during stages of differentiation, we show that bioreactor differentiated clusters were able to retain a considerable yield (∼42%) of potent endocrine cells (Ins^+^Cpep^+^ISL1^+^) within the SC-islets capable of abrogating chemically induced diabetes in immunodeficient mice. Through the final stage of maturation (S6), bioreactor differentiated SC-islets exhibited rationalized endocrine cell composition (monohormonal cells for insulin, glucagon, and somatostatin) and organotypic cytoarchitectural morphology. These observations are correlated and confirmed with decreased cell proliferation (Ki67^+^Ins^+^) and polyhormonal cells (Ins^+^Gcg^+^Som^+^) while presence of relatively lower SC-EC commitment that are insulin expressing, implicating fetal islet development and *in vitro* islet function. Adopting VW® suspension culture approach, the off-target cell generation was comparable and lower than previously reported by Balboa *et. al.,* and Veres. *et.al.,*^7, 15^ within 27-day duration.

Despite fetal maturation phenotype, single cell resolution analysis confirmed absence of higher degree of cellular heterogeneity. Bioreactor-cultured SC-islets attained controlled cellular heterogeneity at S4 progenitors and S6 maturation, against primary donor islets as observed with PDX1, GP2, NKX6.1 CHGA, CPEP, and ISL1 expression. The most intriguing implication of this single cell analysis is an ability of SC-islets to confirm for their proteomic differences for cell composition supporting our observed data for genomic architecture and functional efficiency against primary islets. The levels of cellular maturity at the end of stage-6 was in accordance with advanced optimal 2D planar protocols^6, 7, 15^.

To investigate the endocrine heterogeneity, we used transcriptional gene profiling as a tool to scrutinize the expression of key maturation genes for beta cell development and differentiation leading to enhanced insulin secretion and exocytosis. Our result suggested that 3D culture dynamics of bioreactor facilitated enhanced transcriptional maturity with increased expression of several functional genes of beta cell maturity, glucose sensing and exocytosis. An accelerated SC-islet maturation under suspension culture delivered transcriptional control favouring glycolysis or TCA pathways, at least partly, for robust endocrine maturation and glucose-dependent insulin secretion response, similar to adult primary islets.

VW^®^-based bioreactor enables positive outcomes in preserving cell loss, improving islet yield, and enhancing metabolic, transcriptional, and functionally maturity of cells, *in vitro* and *in vivo*. Furthermore, transcriptomic analysis of the endocrine graft 120 days post-transplantation demonstrated the upregulation of key islet maturation genes including *G6PC2, PCSK1,* and *GLP1R* with further maturation after engraftment. Furthermore, zinc transporter *SLC30A8, ABCC8*, alpha cell maturation marker *ARX* were upregulated while *SLC18A1* expression decreased, suggesting an evolving commitment into β-cell differentiation and maturation. Glucose dependent increased human C-peptide secretion after transplantation showed resetting of glycemia to human levels, and *in vivo* graft function, predominantly due to transcriptional maturation post implantation. We also detected glucose sensing and concentration dependent insulin secretion capacity in graft cells particularly after *in vivo* maturation, consistent with its role in postnatal maturation of fetal islets in mouse pancreatic development. Our findings for functional metabolic, and transcriptomic maturity are in accordance with findings stated by Balboa *et. al.,* and Veres. *et.al.,*^7, 15^ within 27-day duration.

In summary, we present a complete and defined 3D suspension culture protocol for the reliable generation of functional SC-islets, although resembling fetal islets *in vitro* and mature *in vivo* post transplantation similar to adult donor islets. Our findings endorse the VW complete 3D approach an efficient means to validate SC-islet products against “gold-standard” adult primary donor islets. Our protocol offers several advantages. First, the bioreactor system allows for precise control of environmental parameters such as oxygen and nutrient levels, pH, and mechanical stimulation. This can promote cell differentiation and maturation by mimicking the dynamic and heterogenous microenvironment of the pancreas. Second, the bioreactor system has the potential to show further scale up of functional SC-islets for therapeutic cell-mass in humans, which will become critical for future diabetes cell therapies. Third, the system will allow for the optimization of differentiation protocols by monitoring cell composition, quality control parameter for release criteria and genomic regulation with direct integration of automation, robotics and artificial intelligence.

## CONCLUSION

We herein developed a complete 3D suspension culture protocol that obviates a need for 2D plating in the efficient generation of iPSC-derived human islet-like clusters using VW^®^ bioreactor technology. Our study shows advancement and promise for improved islet cell manufacturing and scale up to develop clinical-grade islet cell therapy product for diabetes. Future research will advance the utility of these bioreactors for GMP-grade SC-islet manufacturing. Ongoing work will further optimize these protocols for fluid dynamics, reduced mechanical sheer stress during scale-up, and potentially for additional methods to facilitate targeted removal of unwanted off-target cell populations. Ultimately, the technology has considerable potential to accelerate generation of autologous stem cell-derived islets cell therapies for the treatment of all forms of diabetes without the need for antirejection drugs.

## MATERIALS AND METHODS

### Experimental model and subject details

All procedures and protocols have been reviewed and approved by the Stem Cell Oversight Committee (SCOC), Canada and the University of Alberta Institutional Health Research Ethics Board (PRO00084032). All animal protocols were conducted in accordance with the Canadian Council on Animal Care Guidelines and Policies with approval from the Animal Care and Use Committee (Health Sciences) (protocol number-AUP00000331) for the University of Alberta. Euthanasia was performed by filling the euthanasia chamber at a rate of 25% CO_2_ chamber volume per minute to cause the least amount of distress to rodents. Patients recruited in this study as blood sample donors provided written consent for use of tissue, cell reprogramming, and result disclosure. All experiments were planned *a priori* and completed in biological and technical triplicates based on standard experimental procedures without exclusion of experimental groups. Randomization was not performed but throughout experimental procedures the scientist performing analysis was blinded to the group allocation of samples including grafts after recovery. Other confounders were not controlled for.

### Cell culture

Cell culture was performed in a Class-II biocontainment compliant lab with manipulation of cells in a sterile environment using a biosafety cabinet with high efficiency particulate air filtration. Cells were maintained at 37°C with 5% CO_2_ within humified incubators.

### Generation, maintenance, and expansion of human induced pluripotent stem cell lines

Two human iPSC lines generated from peripheral blood mononuclear cells (PBMCs) from healthy donors (patient demographics in Supplementary Material Table S1) were used in this study. Donor blood (20 mL) was collected in a sterile fashion into BD vacutainer spray coated K2EDTA tubes (Fisher Scientific, cat. 13-680-61) and PBMCs were isolated using density gradient centrifugation on Histopaque and expanded by using StemPro-34 Serum Free Complete Media (Gibco, cat. A14509) supplemented with cytokines (10 ng/mL IL3, IL6, SCF and FLT3; R&D Systems, cat. 203-GMP, 206-IL, 7466-SC, and 308E-GMP respectively) for four days to achieve satisfactory cell numbers prior to iPSC reprogramming with Sendai virus. iPSCs were generated using the CytoTune iPS 2.0 Sendai Reprogramming Kit (Thermo Fisher, cat. A16517). Between days 15-20, a minimum of 10-12 individual colonies (referred as clones hereafter) were manually isolated by handpicking under 10X phase objective (using ECHO inverted Rebel microscope and ECHO image acquisition application on 10.1in iOS device (ECHO)) and subcultured as unique iPSC clone cell lines. Each clonal cell line was scrutinized for routine pluripotent stem cell quality control criteria (ALP staining, phenotype, genetic analysis, and genomic integrity) with the best clone used to establish a single and normal iPSC cell line for use in this study.

iPSC lines were cultured on 60-mm rhVTN coated plates in StemFlex media (Stem Cell Technologies, cat. A3349401) and passaged using CTS EDTA Versene Solution (Fisher Scientific, cat. A4239101) supplemented with 2 μl/mL Rho-kinase inhibitor (RockI; Y-27632 STEMCELL Technologies, cat. 72304). To prepare the differentiation experiments, near-confluent plates of iPSCs were dissociated and seeded on 150-mm Geltrex coated plates in StemFlex supplemented with 2 μl/mL ROCKi at a density of ∼0.06 million cells cm^−2^ and expanded for 3 days to achieve 80 to 90% confluency prior to differentiation.

For expansion in Vertical Wheel (VW) Bioreactors, 2 million iPSCs acquired from 60-mm rhVTN coated plates were adapted to suspension culture in 0.1 L VW Bioreactors (PBS Biotech, cat. FA-0.1-D-001) for 3 passages before any experimental usage. At 4^th^ passage onwards, upon passaging, cells were counted, and 10 million live cells were seeded into 0.5 L VW Bioreactors (PBS Biotech, cat. no. FA-0.5-D-001) in 250 mL of StemFlex media with RockI (2 µL/mL) (day 1) with constant rotational speeds of 60 rotations per minute (rpm). After 24hr, clustered iPSCs were then supplemented with an additional 250 mL of StemFlex media without RockI. On days 4 clusters were allowed to settle down by gravity for 10 minutes inside the incubator; the upper 250 mL of StemFlex was removed and 250 mL of fresh pre-warmed StemFlex was replenished in the reactor. On day 6, clusters were ready for differentiation.

### Differentiation of induced pluripotent stem cell lines

For planar differentiation, iPSCsh were seeded into geltrex coated plates in StemFlex media for expansion and the medium was changed to S1 medium 3 days post-seeding. For suspension differentiation, iPSCs were seeded into a 0.5 L VW Bioreactor for expansion and the medium was changed to S1 medium 5 days post-seeding; differentiation was performed at 60 rpm. In both cases, differentiation was carried out using a six-stage protocol modified from key publications ^8, 21^. Complete media formulations for planar and suspension differentiation are available in the Supplementary Material Table S2. For planar differentiation, on day 12, the planar pancreatic progenitors were lifted using 10 min Accutase treatment (Thermo Fisher Scientific, cat. A11105-01) supplemented with 2 μl/mL ROCKi and seeded into microwells (6-well AggreWell^TM^ 400 plates, Stem Cell Technologies, cat. 34460) at a density of 800–1,000 cells per microwell using the manufacturer’s recommended protocol. After 24 hours, the clusters were transferred from the microwells to suspension culture on 0.1 L VW Bioreactors at 60 rpm. Media changes were performed daily on S1 and every 2 days thereafter (refer to Supplementary Material Table S2).

### Bioreactor imaging

Photomicrographs of islet-like clusters developing and differentiating in VW® was captured with Apple iPhone 7 Plus using 12-megapixel wide angle camera-ISO 25, 28mm, 0 ev, *f* 1.8 and 1/60s shutter speed. All images were captured as 3024 x 4032 pixels.

### Isolation of primary adult islets

Human islets were obtained from The Alberta Diabetes Institute Islet Core (University of Alberta, Canada) after donor family research consent, and after Health Research Ethics Board review (Pro00001620). Islet donor characteristics are listed in Supplementary Material Table S3.

### Alkaline phosphatase staining

Following iPSC clone selection, individual clones were grown on geltrex coated 24×24 mm glass coverslips in 6 well plates, fixed in 4% paraformaldehyde (PFA) for 10 minutes and incubated with ALP-substrate staining solution (Abcam, cat. Ab242287) for 20 minutes in the dark at RT as per manufacturer recommendations. Positively stained colony images were visualized and analyzed with ECHO Rebel inverted microscope (ECHO).

### Genetic analysis

Whole genomic DNA (gDNA) extraction was performed by lysing a maximum of 5 x 10^6^ cells for 18-24hr at 55°C 487.5 µL TENS buffer (10 mM Tris-HCl (Sigma, cat. T3253) pH 8.0, 25 mM EDTA (Sigma, cat. 324506) pH 7.5, 100 mM NaCl (Sigma, cat. S1679), 0.5% SDS (Sigma, cat. 71736)) with 12.5 µL proteinase K (20 mg/mL; Sigma, cat. 70663-4). Following cell lysis, proteins were precipitated with 250 µL of 6 M NaCl and centrifuged for 5 min at 12,0000 x g. The supernatant was recovered, and gDNA was precipitated with 900 µL isopropanol followed by centrifugation at 12,000xg for 10 min. The pellet was collected and washed with cold 70% ETOH and then allowed to dry. Pelleted gDNA was resuspended in 50 µL TE buffer (10 mM Tris-HCl pH 8.0, 1 mM EDTA pH 8.0) and purity was assessed using the Multiskan SkyHigh Microplate Spectrophotometer and µdrop plate (Thermo Scientific). Sample purity was assessed by measuring the 260/280 nm and 260/230 nm absorption and only samples achieving 1.7-2.0 and 2.0-2.2 respectively were further evaluated.

For genetic analysis, reactions were set up in 96 wells plates using the hPSC Genetic Analysis Kit (STEMCELL technologies, cat. 07550) as per manufacturer instructions with triplicates for each sample. qPCR was performed using the StepOnePlus Real-Time PCR System (Fisher Scientific, cat. 4376600) and gDNA was amplified as per Supplementary Material Table S4. Samples were analyzed using STEMCELL Technologies’ genetic analysis app (https://shiny.stemcell.com/ShinyApps/psc_genetic_analysis_app/), which uses chromosome 4p as reference for data normalization and calculates copy number as 2^(-ΔΔCT)^ x 2, which enables the visualization of copy number for the specific chromosomal regions of each of the tested samples.

### Immunohistochemistry and image analysis

iPSCs and samples differentiated on planar conditions were grown and differentiated on coverslips coated with geltrex and fixed in 4% PFA for 20 minutes at RT. Samples differentiated in suspension conditions and explanted SC-islet grafts were fixed overnight in 4% PFA and embedded in paraffin. Sections (5 μm) were deparaffinized and rehydrated and subjected to antigen retrieval using citrate buffer (0.0126 M citric acid, Sigma cat. C-0759; 0.0874 M sodium citrate, Sigma, cat. S-4641; pH 6.0) for a total of 20 minutes. Coverslips and slides were then blocked and permeabilized with 5% normal donkey serum (Sigma, cat. S30-M) in FoxP3 permeabilization buffer (Biolegend, cat. 421402) for 1 h at room temperature (RT) and incubated with primary antibodies diluted in permeabilization buffer overnight at 4 °C. Secondary antibodies were diluted similarly and incubated for 1 h at RT and DAPI (Sigma, D1306) staining for 4 minutes at RT was used for nuclear visualization. Antibodies are listed in Supplementary Material Table S5. Slides were visualized using the Zeiss Observer Z1 inverted fluorescence motorized microscope and images were processed using the Zen2 Blue Edition v.2 (Zeiss) and analyzed using QuPath ^36^.

### Flow cytometry

Pancreatic stage-specific cells (Stage1-5) and stage 6 SC-islets were dissociated with pre-warmed accutase for 10 min in a 37 °C water bath, sound down and resuspended in 1 mL PBS. Cells were strained through a 40 µm strainer, fixed with 4% PFA for 20 minutes at RT and permeabilized using Cytofix/Cytoperm (BD Biosciences, catalog no. 554714) for 20 min on ice followed by twice washes with 1x Perm/Wash buffer (BD Biosciences, catalog no. 554714). Primary antibodies were incubated for 1 h on ice, or overnight at 4 °C when required, and secondary antibodies for 30 minutes according to the dilutions in Supplementary Material Table S5; cells were resuspended in fluorescence – activated cell sorting buffer (2% FCS, 2 mM EDTA in DPBS). Data were acquired using the CytoFLEX S flow cytometer and analysed using the CytExpert software (Beckman Coulter), FlowJo v.10 (BD Biosciences) and Cytobank v10.0 premium subscription (Beckman Coulter). FlowSOM and tSNE analysis were performed using Cytobank v10.0 ^37^, which uses self-organizing maps for dimensional reduction visualization of flow cytometry data.

### Gene expression

Cells were lysed with 350 μL RLT buffer (Qiagen, cat. 79216) and frozen at - 80 °C until RNA extraction. 30 mg of grafts were lysed in RLT buffer followed by sonication and frozen at -80 °C until RNA extraction. Suspension of lysed cells or tissues in RLT buffer was defrosted and cells were disrupted and homogenized using the QIAshredder system (Qiagen, cat. 79656) and total RNA was then extracted with the RNeasy Mini Kit (Qiagen, cat. 74104) according to the manufacturer’s instructions. Concentration and purity of the isolated RNA samples was evaluated using spectrophotometry with the Multiskan SkyHigh Microplate Spectrophotometer and µdrop plate (Thermo Fisher, cat. A51119600DPC) by assessing the 260/280 nm and 260/230 nm absorption of samples. Samples were then stored at -80 ⁰C until needed; RNA was quantified after each defrost. RNA was reverse transcribed using the RevertAid First Strand cDNA Synthesis Kit as per manufacturer guidelines (Thermo Fisher, cat. K1621). Complement DNA (cDNA) was stored at -20 ⁰C until required for PCR.

To quantitatively study gene expression, custom designed gene TaqMan Low Density Array Cards were procured and used as per manufacturer instructions (Fisher Scientific cat. 4342253); gene array set up is described in Supplementary Material Table S6. Briefly, 500 ng of cDNA were combined into a solution with 55 μL of nuclease free water and 55 μL TaqMan Universal PCR Master Mix (Thermo Fisher cat. 4305719). The combined solution was loaded into the gene array cards, centrifuged, and processed using FAST-384 well array program via the QuantStudio 12K Flex Real-Time PCR system as per the manufacture recommendations. Data was then analyzed as above and represented as a heat map and/or 2(-ΔΔCT) and/or volcano plots using GrapBio ^38^ and https://www.bioinformatics.com.cn/en, a free online platforms for data analysis and visualization.

### Glucose stimulated insulin secretion

Static tests of insulin secretion were carried out in 1.5 mL tubes. A total of 30-50 SC-islets were handpicked and equilibrated in Krebs-Ringer buffer (KRB) with 2.8 mM glucose (G3) for 120 min, and then subjected to sequential 30-min incubations of G3, 16.8 mM glucose (G17), 100 nM Exendin-4 (Sigma, cat. E7144) and 30 mM KCl in KRB. After the tests, the SC-islets and supernatant were collected, and the insulin and DNA contents were analyzed. Dynamic tests of insulin secretion were carried out using a perifusion apparatus (Brandel Suprafusion SF-06) with a flow rate of 0.1 ml min–1, and sampling every 1 min. A total of 50 handpicked SC-islets were used for each test. The SC-islets were equilibrated in G3 for 120 min prior to sample collection. Insulin content of secretion fractions and SC-islet lysates was analyzed with enzyme-linked immunosorbent assay (ELISA) (Mercodia, cat. 10-1132-01).

### Diabetes induction and SC-islet transplantation

Five days before transplantation, diabetes was induced by intraperitoneal (IP) injection of 175 mg/kg of streptozotocin (STZ; Sigma-Aldrich, ON, Canada) in acetate buffer, pH 4.5 (Sigma-Aldrich, ON, Canada). Immunocompromised SCID beige mice (10 to 16-week-old) mice were considered diabetic following a non-fasting blood glucose measurement ≥15.0 mmol/L on two consecutive days. Pancreatic progenitors or SC-islets were separated into tubes with 1.5 million beta-like cells (Nkx6.1^+^ Isl-1^+^ C-peptide^+^), ∼2,000 human islet equivalents, and transplanted under the kidney capsule as previously described ^40^. Institutional guidelines for perioperative care, anesthesia and pain management were followed.

### Evaluation of SC-islet graft function

Glycemic control was assessed using non-fasting blood glucose measurements (mmol/L) thrice weekly after transplantation using a portable glucometer (OneTouch Ultra 2, LifeScan, Canada). Diabetes reversal was defined as two consecutive readings <11.1 mmol/L. SC-islet grafts were retrieved by total nephrectomy three and/or six months after transplantation. Retrieved grafts were characterized using IHC, flow cytometry and/or RT-qPCR. For IHC, complete or partial grafts were fixed overnight in 4% PFA and embedded in paraffin as previously described. For flow cytometry analysis and glucose stimulated c-peptide response, grafts were digested with 2mg/ ml collagenase Type V (Sigma, cat. 9623) for 10 min in a 37 °C water bath with shaking every 2.5 min. Single cells were fixed in PFA for flow cytometry analysis or clustered for 24 hours in 6-well AggreWell 400 plates a density of 800–1,000 cells per microwell in stage 6 media for glucose stimulated c-peptide response assay. Lastly, for RT-qPCR, up to 30 mg of grafts were stored in RLT buffer and frozen at -80 °C.

Furthermore, glucose tolerance tests were conducted at 8, 12 and 16 weeks after transplant, as a means to further assess metabolic capacity in response to a glucose bolus, mimicking postprandial stimulus. Animals were fasted overnight before receiving an intraperitoneal glucose bolus (3 g/kg). Blood glucose levels were monitored at 0, 15, 30, 60, 90 and 120 min after injection, allowing for area under the curve (AUC)-blood glucose to be calculated and analyzed between transplant groups. C-peptide content at 0 and 60 minutes after injection were analyzed with enzyme-linked immunosorbent assay (ELISA) (Alpco, cat. 80-CPTHU-CH01).

### Data collection and statistical methods

Morphological data represents population-wide observations from independent differentiation experiments. viSNE and FlowSOM dimensional reduction visualization and insulin secretion and transcriptomic data represents samples of independent SC-islet differentiation experiments or islet donors. *In vivo* data is derived from independent animals. Statistical methods used are represented in each Figure legends. All statistical analysis was performed using one-way analysis of variance (ANOVA), multiple comparison using Turkey correction, unpaired parametric t-test, and two-way ANOVA with multiple comparison using Šidák correction with 95% confidence interval. All data are presented as either median with 25-75 percentile interval or mean±sem.

## Acknowledgments

AMJS is supported through a Canada Research Chair (Tier 1) in Regenerative Medicine and Transplant Surgery, and through grant support from the Canadian Stem Cell Network, Canadian Institute of Health Research, Juvenile Diabetes Research Foundation, Diabetes Canada, Canadian Donation and Transplant Research Program (CDTRP), Alberta Diabetes Foundation and Diabetes Research Institute Foundation of Canada (DRIFCan). Braulio A. Marfil– Garza is supported by the CHRISTUS Excellence and Innovation Center. We thank Dr. Jean Buteau for providing access to Olympus slide scanning imaging station.

## Author contributions

Following authors have contributed to the preparation of this research article.

Conceptualization: ND, MBL, AMJS

Methodology: ND, MBL, RP, PMD, GJ

Investigation: ND, MBL, NCG, ITJ, RP, BMG, KV, SS, HR, JM

Visualization: ND, MBL, NCG, PMD, AMJS

Funding acquisition: ND, AMJS

Project administration: ND, RP, AMJS

Supervision: ND, AMJS

Writing – original draft: ND, NCG, AMJS

Writing – review & editing: ND, NCG, RP, AMJS

## Data and materials availability

All data are available in the main text or the supplementary materials. Source and extended data are provided with the paper.

## List of Supplementary Materials

Materials and Methods; Supplementary data file S1-S3; Table S1-S7; Extended Data file-1-8; and Extended Source Data Table-1

