## Supplementary figures 1-3 for "Complete Suspension Differentiation of Human Pluripotent Stem Cells into Pancreatic Islets Using Vertical Wheel^®^ Bioreactors"

**Sup. Fig.1: Generation and characterization of human induced pluripotent stem cells from peripheral blood mononuclear cells.**

**Sup. Fig.2: Flowcytometry characterization of stage specific cell composition generated in suspension bioreactors from time of origin.**

**Sup. Fig.3: Flowcytometry and histological characterization and quantification for graft-harvested cell composition.**

### Supplementary data-1

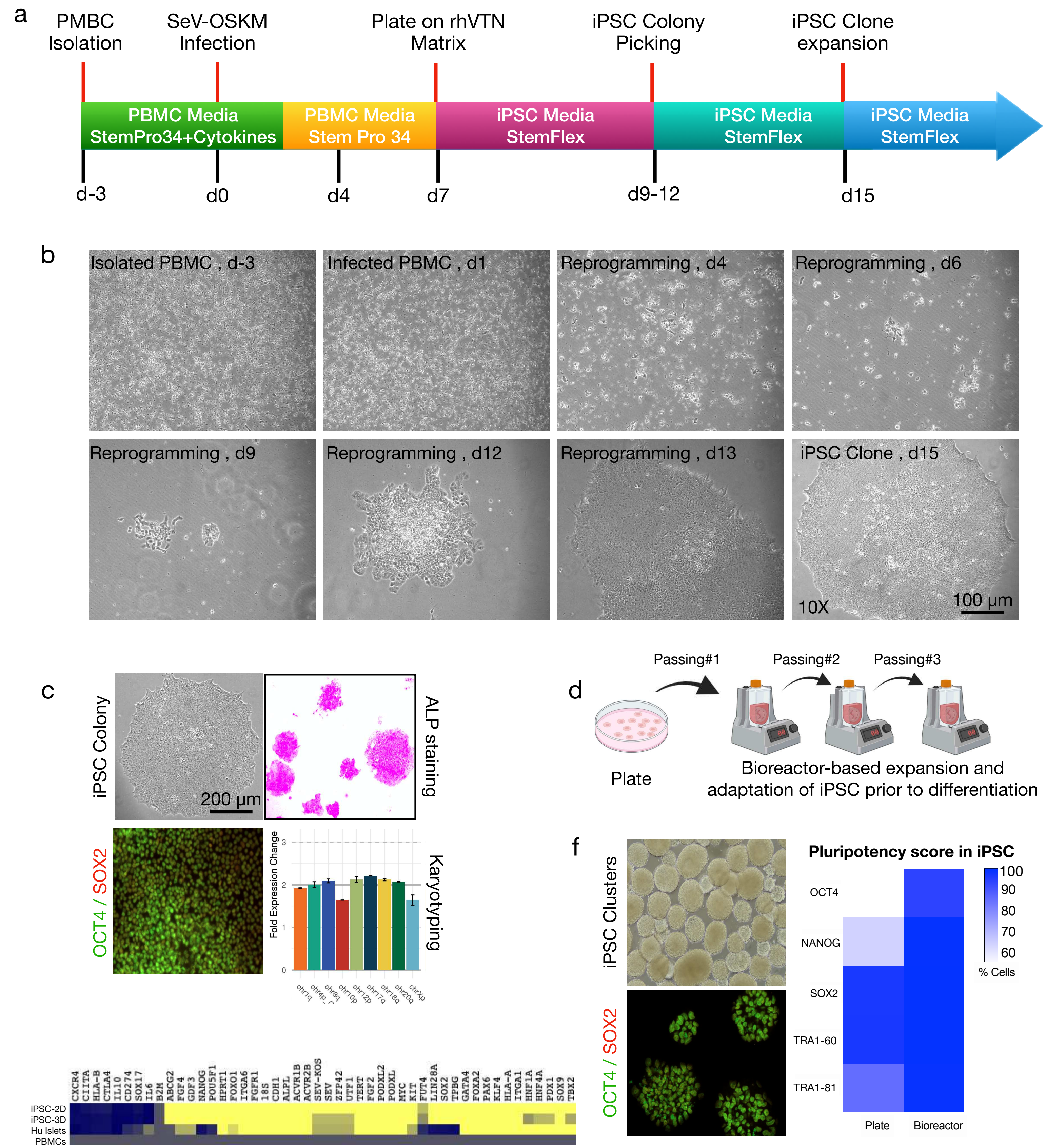

### Supplementary data-2



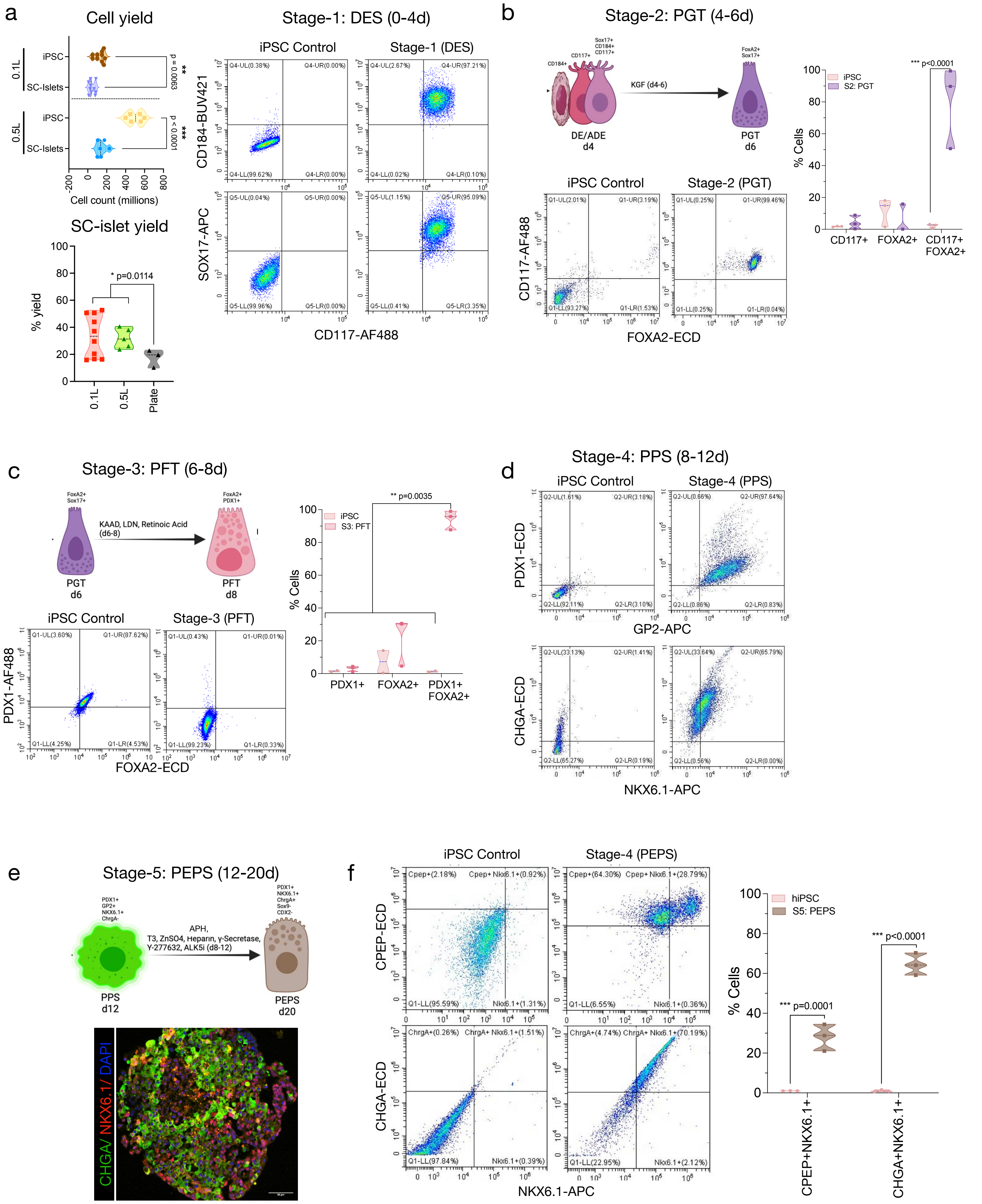

### Supplementary data-3

Dadheech *et al.*, 2023

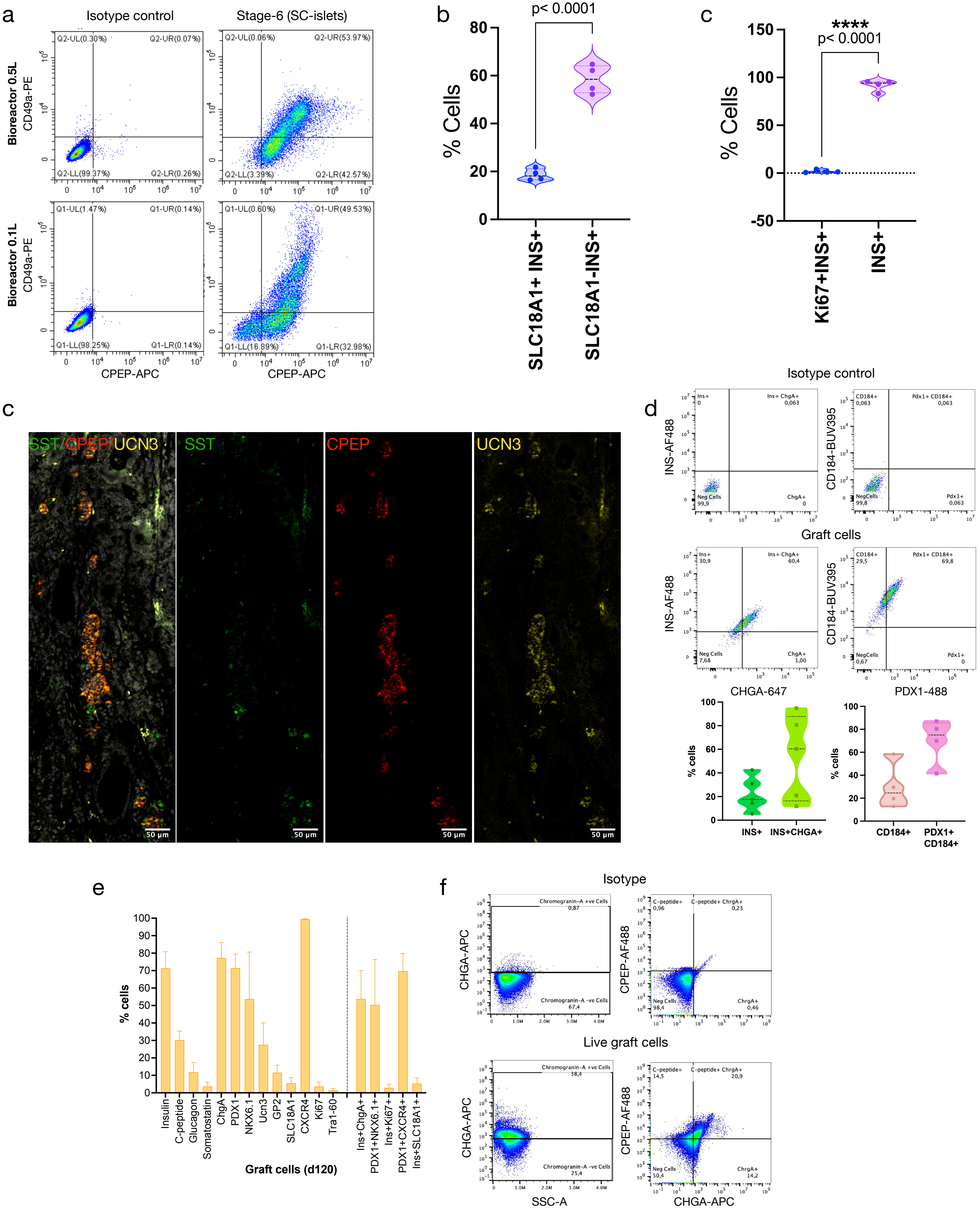
