## Extended Source Data 1-9 for "Complete Suspension Differentiation of Human Pluripotent Stem Cells into Pancreatic Islets Using Vertical Wheel^®^ Bioreactors"

**Extended data-1: Representative flowcytometry characterization of stage-4 cell composition: a,** Flowcytometry plots of stage-4 cells characterized for pancreatic progenitor markers in dispersed single cells from stage-4 clusters generated in suspension bioreactors. b, Flowcytometry plots of stage-4 cells characterized for pancreatic progenitor, proliferation, and pluripotency markers in dispersed single cells from adherent stage-4 differentiated in 2D planar protocol.

**Extended data-2: Representative flowcytometry characterization of stage-6 cell composition: a,** Flowcytometry plots of stage-6 cells characterized for islet and endocrine markers in dispersed single cells from S6w1 cells generated in suspension bioreactors.

**Extended data-3: Representative single cell resolution of flowcytometry-based FlowSOM dimensional reductional analysis of stage-4 cells: a,** FlowSOM projections for pancreatic progenitor markers in stem cell-derived S4w0 single cells generated from three patient-derived iPSC lines compared to adult donor pancreatic islets. **b,** the minimal spanning trees of the self-organizing maps representing unsupervised clustering of stage-4 cell composition based on pancreatic progenitor protein expression levels in single cells compared to undifferentiated iPSCs. **c,** Representative stacks of metaclusters identified in viSNE islands from single cells of stage-4 clusters based on pancreatic progenitor protein expression.

**Extended data-4: Representative single cell resolution of flowcytometry-based FlowSOM dimensional reductional analysis of stage-6 cells: a,** FlowSOM projections for beta cell-specific markers in stem cell-derived S6w1 single cells generated from three patient-derived iPSC lines compared to adult donor pancreatic islets and iPSC control. **b,** the minimal spanning trees of the self-organizing maps representing unsupervised clustering of stage-6 cell composition based on islet hormones expression levels in single cells compared to adult donor islets. **c,** Representative stacks of metaclusters identified in viSNE islands from single cells of S6w1 clusters based on human islet protein expression.

**Extended data-5: Representative single cell viSNE projections of S6 flowcytometry data: a,** viSNE projections for quantification of islet-specific markers in S6w1 single cells differentiated with n=3 independent iPSC lines and compared against adult donor islets and iPSC control cells. **b,** Quantification for islet hormonal population projected within the viSNE islands based on protein expression in S6w1 single cells and gated using human donor islets as reference cell composition. Two-way ANOVA and multiple comparison using Šidák correction with 95% confidence interval was performed. All data are presented as mean±sem, $n=3$, $^{**}p<0.001$.

**Extended data-6: Representative single cell viSNE projections of S6 flowcytometry data: a,** viSNE projections for quantification of endocrine progenitor markers in S6w1 single cells differentiated with n=3 independent iPSC lines and compared against adult donor islets and iPSC control cells. **b,** Quantification for endocrine progenitor population projected within the viSNE islands based on protein expression in S6w1 single cells and gated using human donor islets as reference cell composition.

**Extended data-7: Transcriptomic characterization of stage specific cell composition during islet differentiation using TaqMan Low Density Arrays: a,** Volcano plot visualization of SC-islets (S6w1) against iPSC control displaying expression of key upregulated and downregulated genes. **b,** Volcano plot visualization of Human donor islets against iPSC control displaying expression of key upregulated and downregulated genes. **c,** Principal component analysis to measure an association of stage1-6 transcriptomics against adult donor islets. **e,** Quantification for real time gene expression trajectory in time course during stage-1 to stage-6 differentiation in comparison to adult donor islets.

**Extended data-8: Violin plots projection to show transcriptomic maturation of S6w1 islets in comparison to adult islets:** Violin plots for 48 genes encoded in gene arrays to quantify transcriptome level of S6w1 cells against primary donor islets and reference iPSC control cells.

### Extended Data-1

a

Flow cytometry: Stage 4 cell composition under bioreactor differentiation

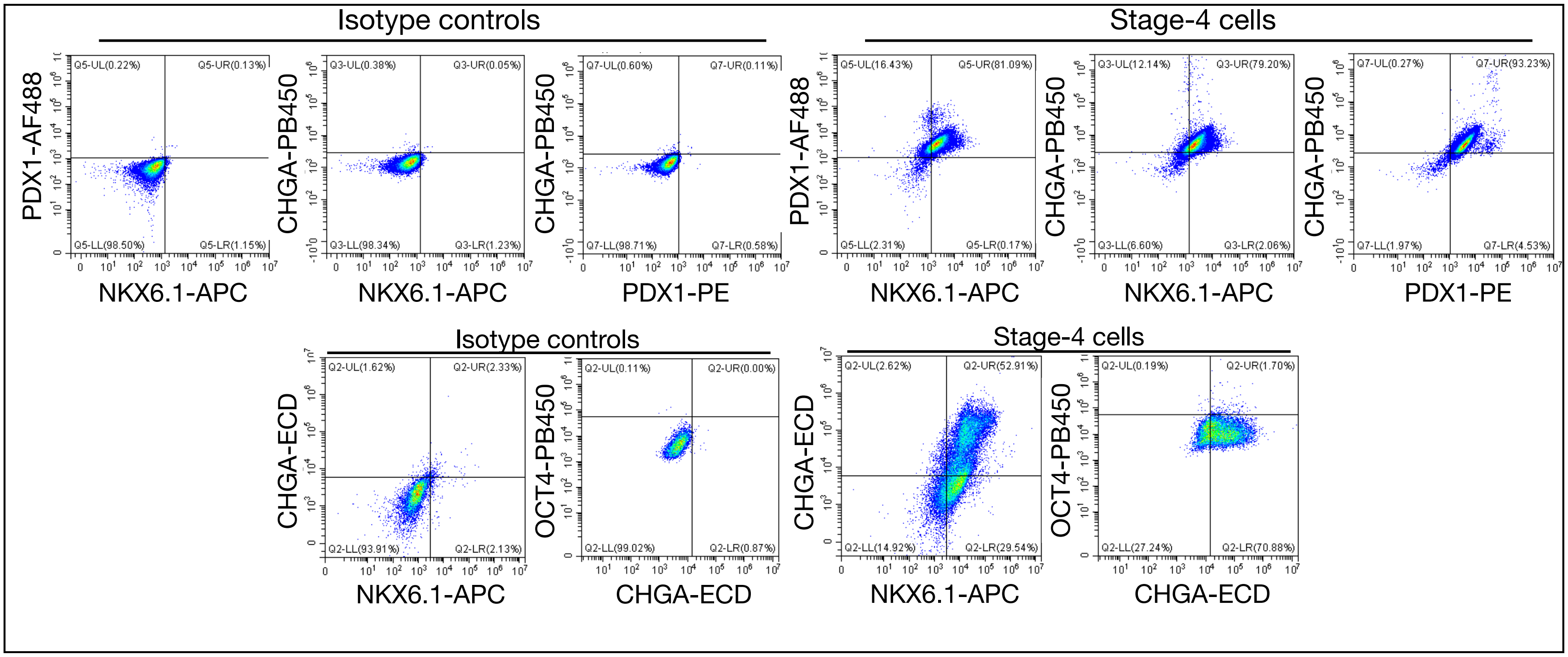


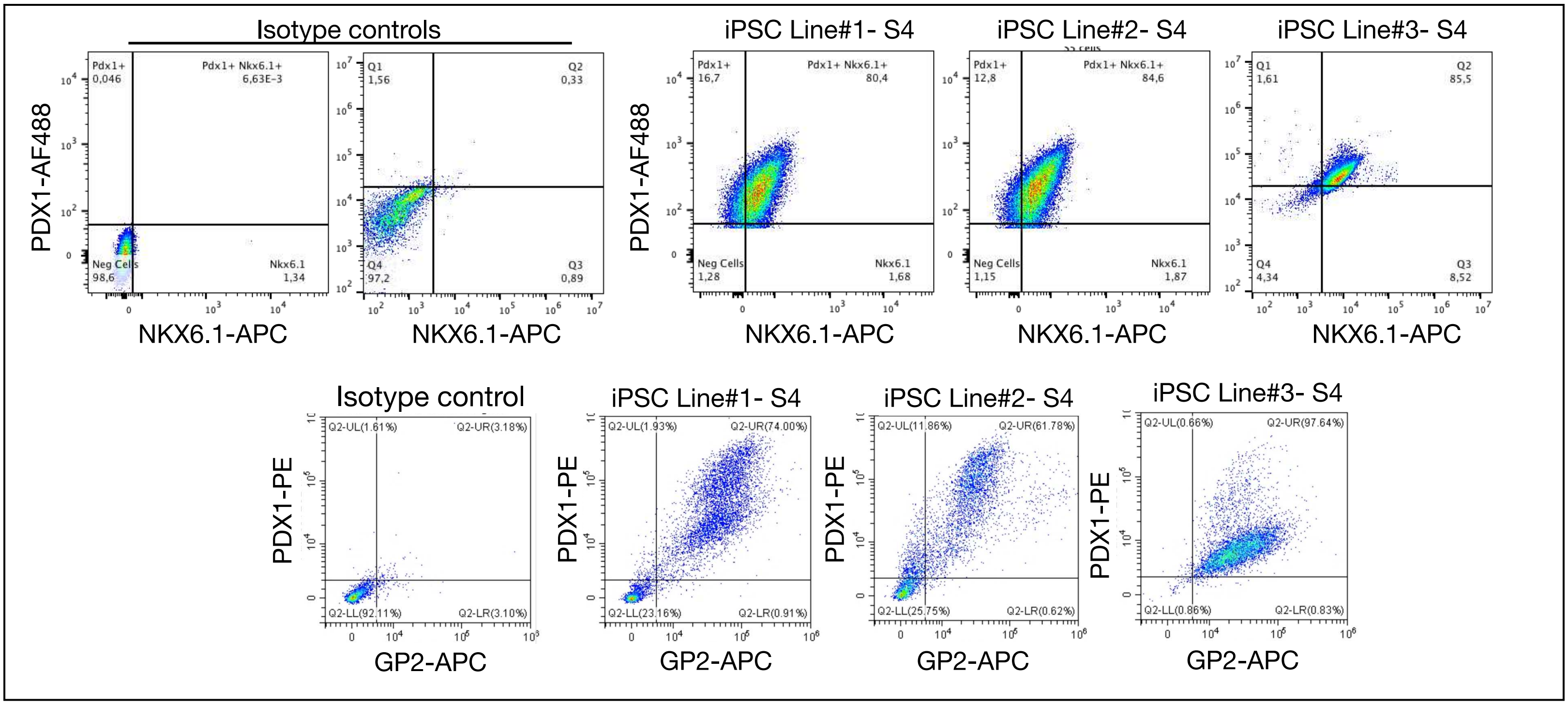

### Extended Data-2

a Flow cytometry: Stage-6 cell composition under bioreactor differentiation

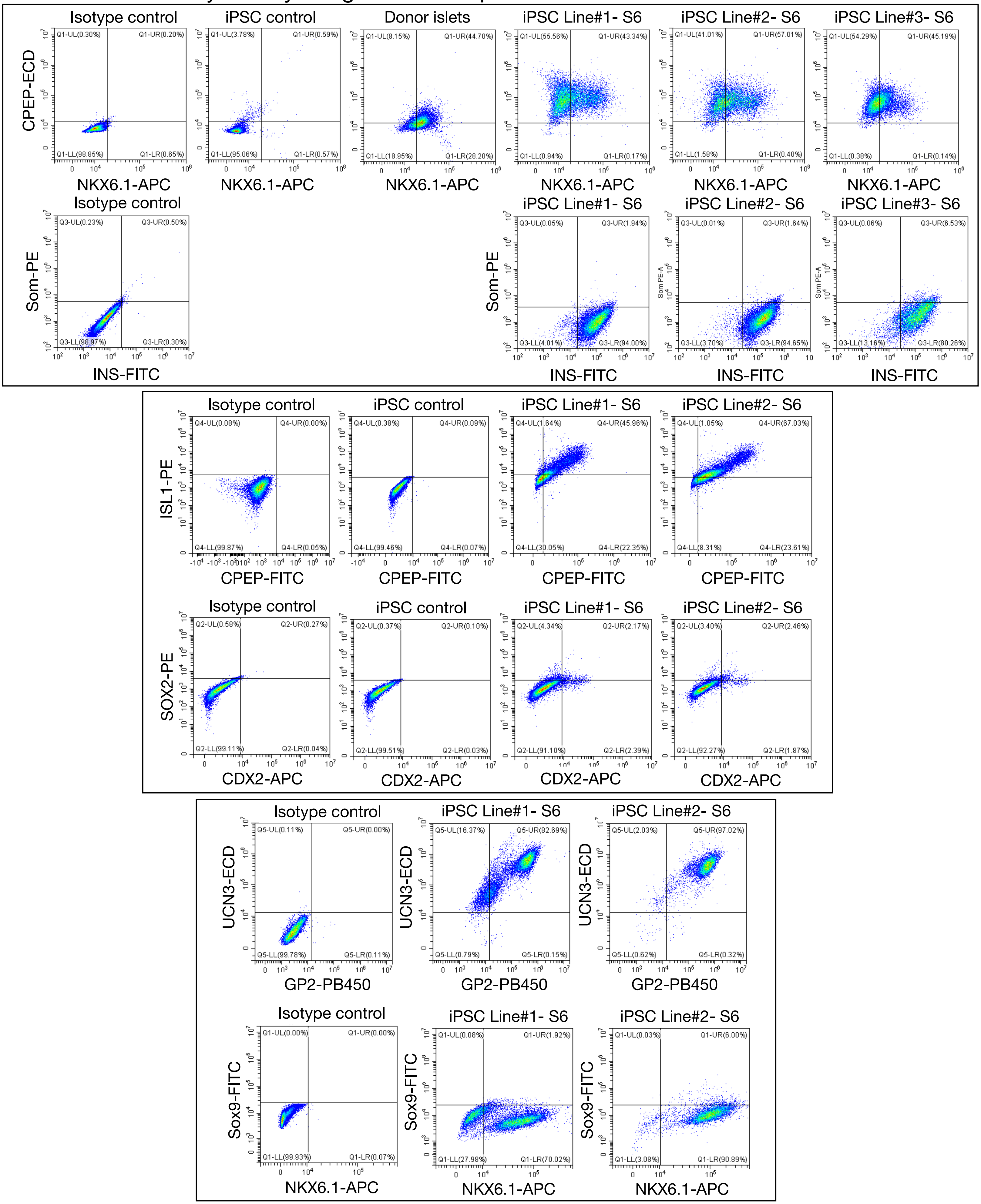

### Extended Data-3

a FlowSOM dimensional reductionality : Stage-4 cells

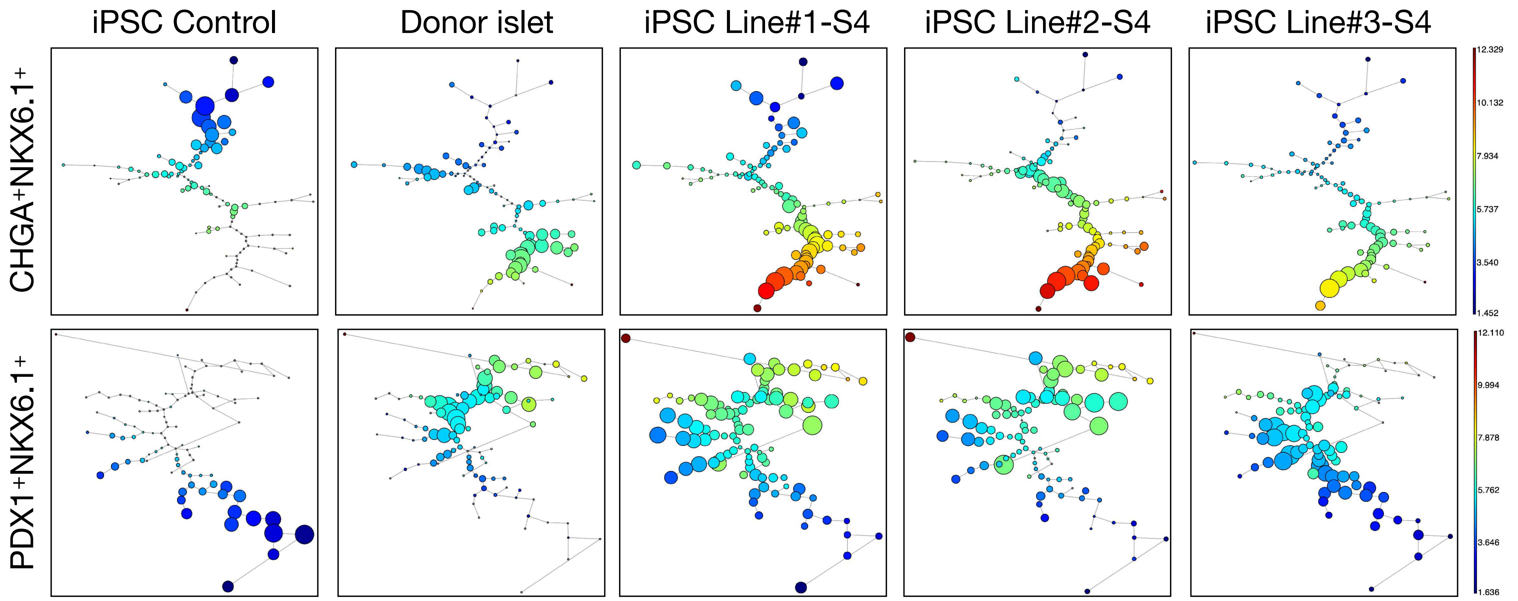


b Star Plots- Self organizing maps : Stage-4 cells

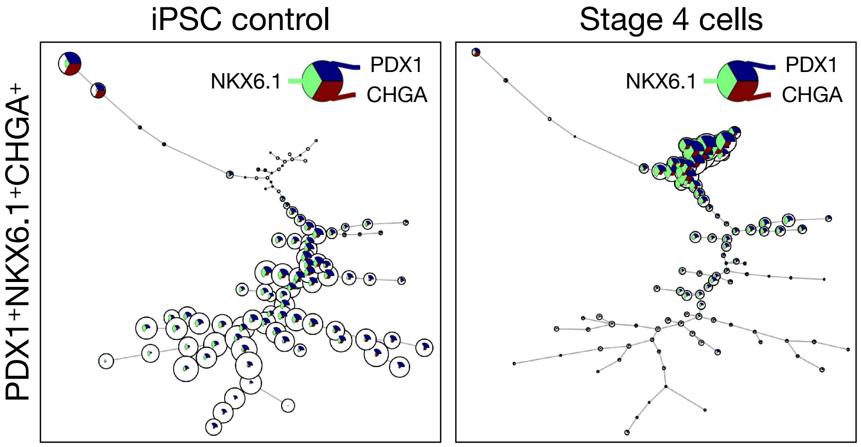


c FlowSOM metaclusters arrangement: S4 cells Vs iPSC

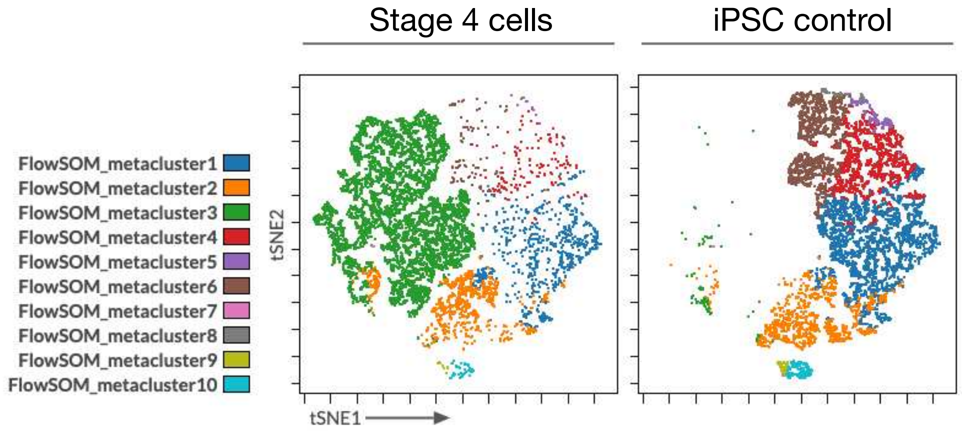

### Extended Data-4

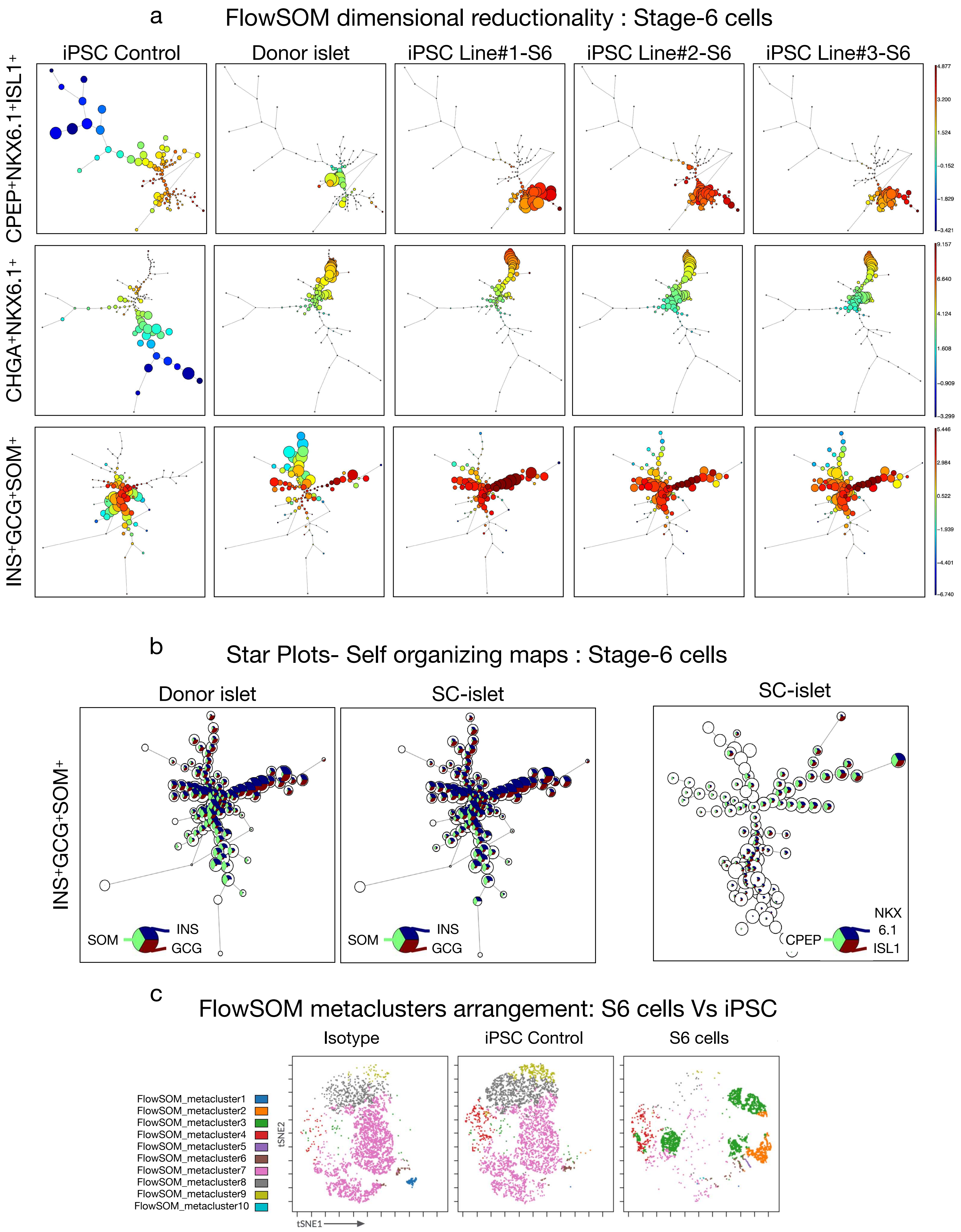

### Extended Data-5

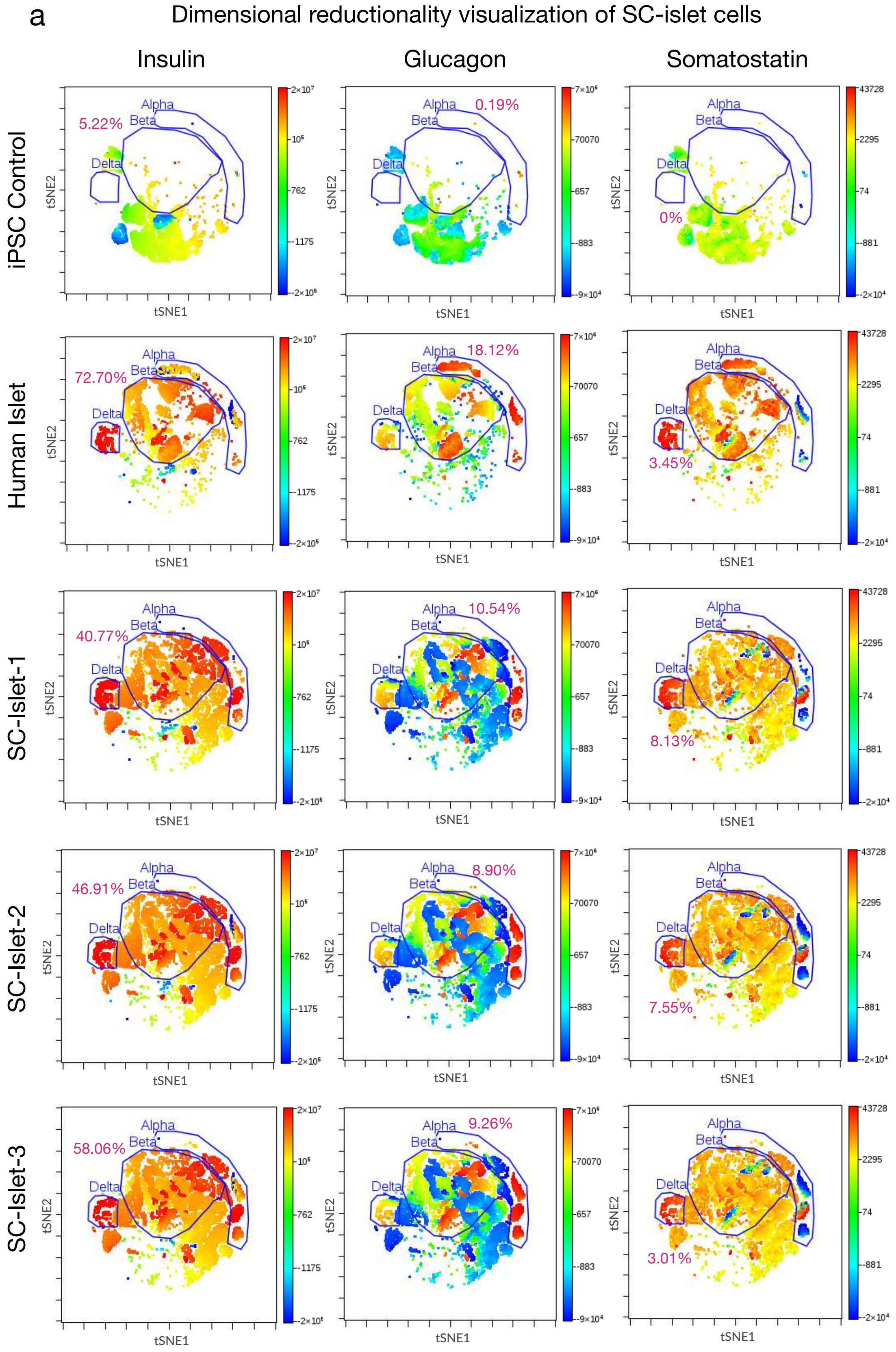


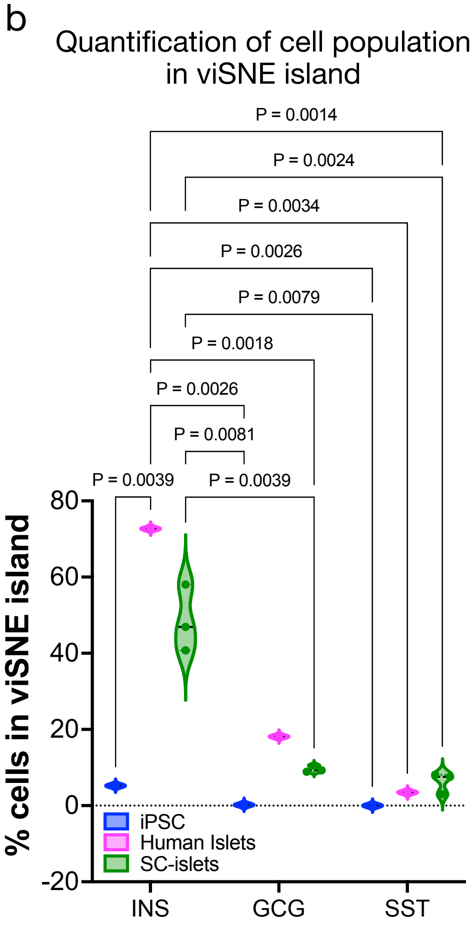

### Extended Data-6

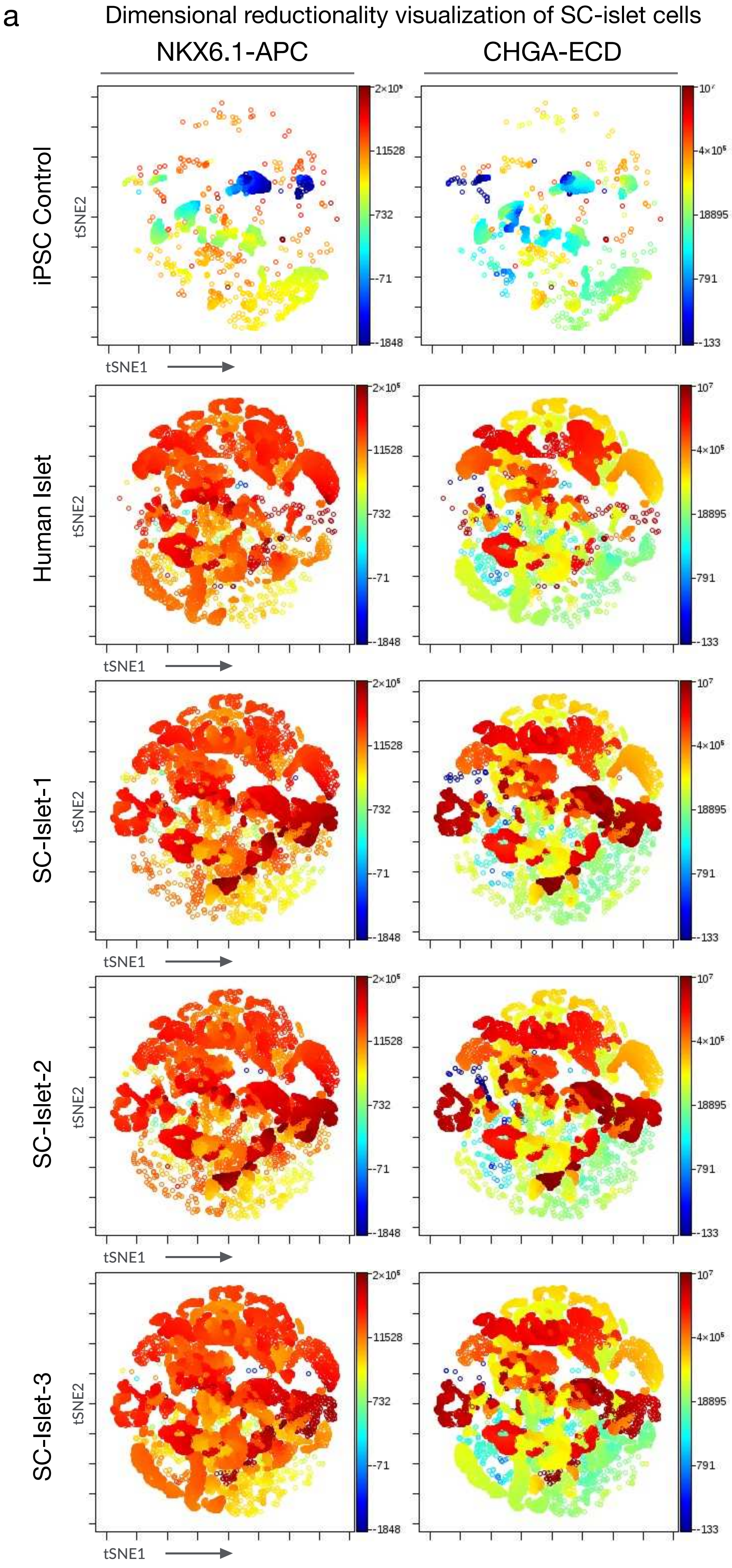


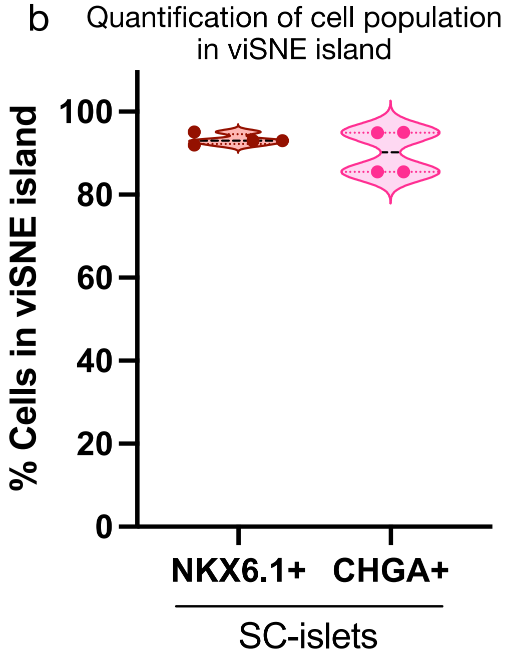

### Extended Data-7

a Volcano plot: Human donor islet vs SC-islet (S6)

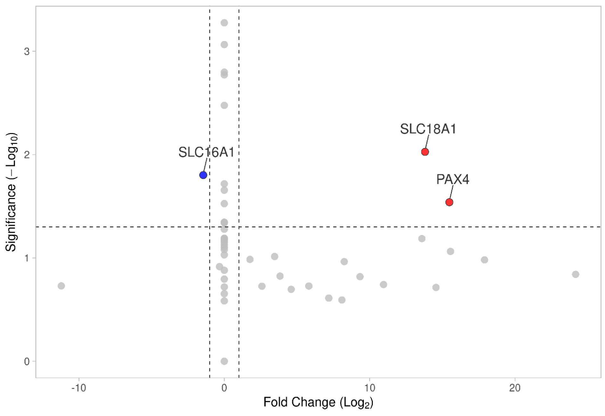


b Volcano plot: Human donor islet vs iPSC

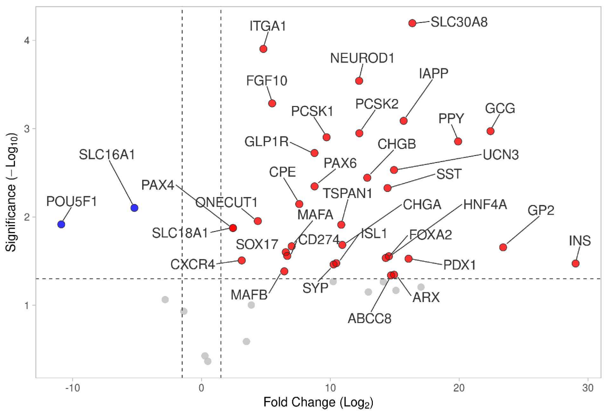


c Principal component analysis: S1-S6 cells

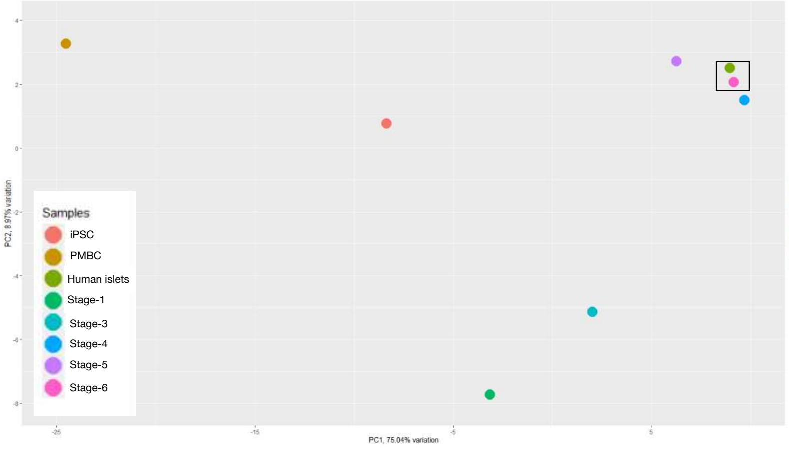


d Stage-specific real-time gene expression profiling

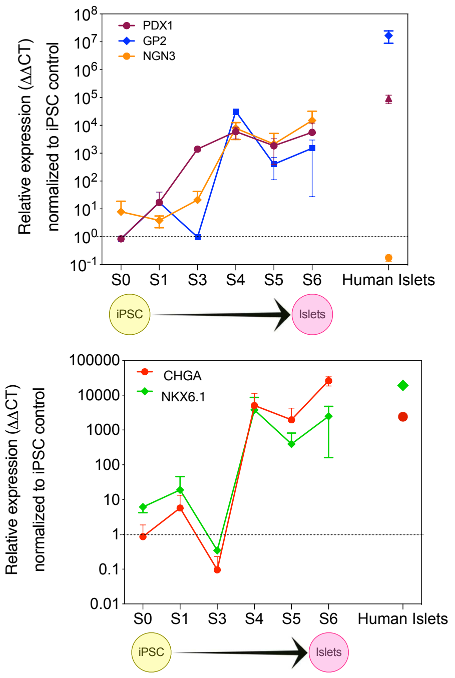

### Extended Data-8

Gene expression profile of 48 islet cell differentiation genes

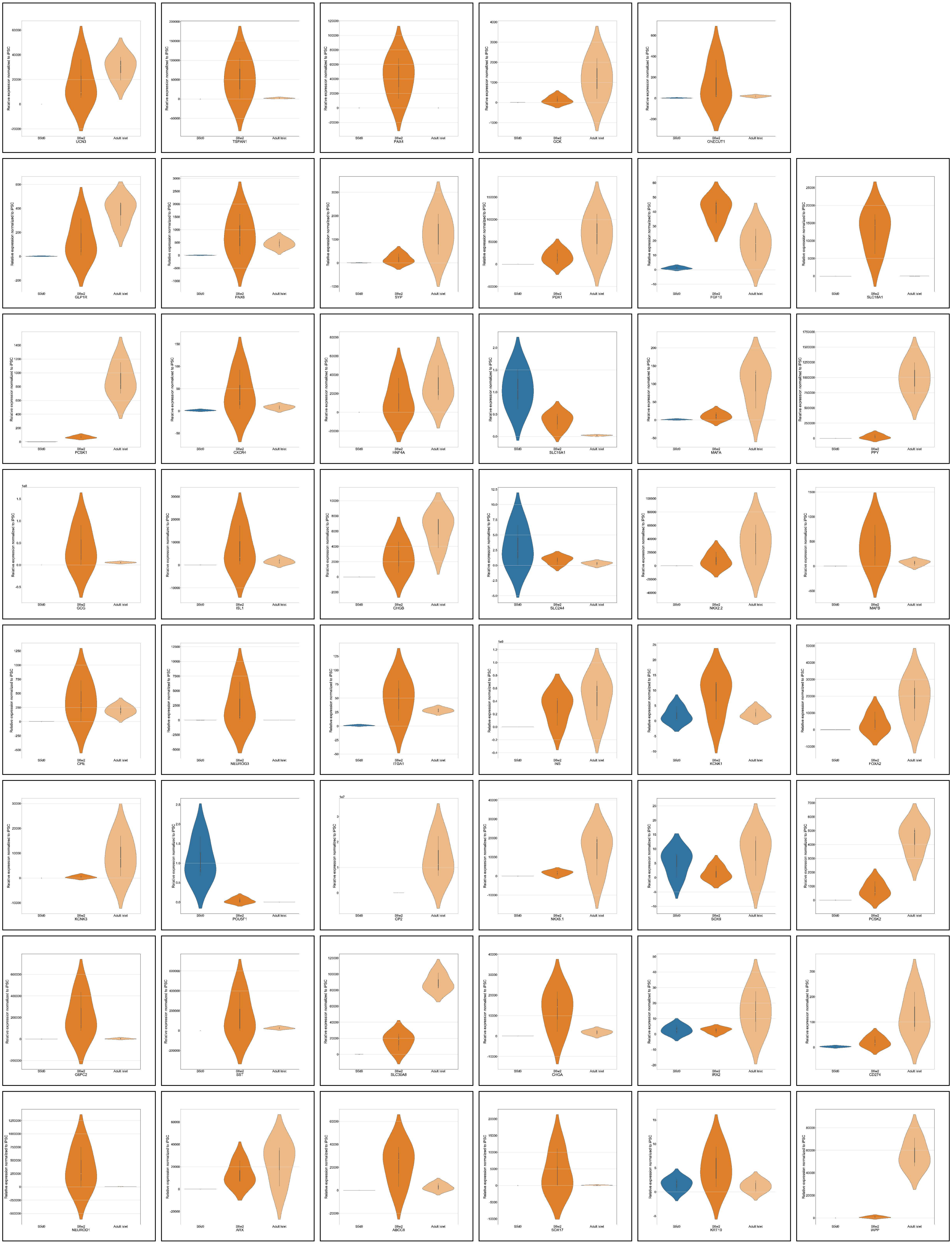

### Extended Data- Table 1

Gene Expression: Volcano Plots Statistics

Human Donor Cadaveric Islets versus Human Induced Pluripotent Stem Cells

### VolcaNoseR - Exploring volcano plots

| Name | Change | Fold change (log2) | Significance | Manhattan distance |
|---|---|---|---|---|
| INS | Unchanged | 29.037739 | 1.47301215161794 | 30.5107511516179 |
| GCG | Increased | 22.4330126666667 | 2.97127893883463 | 25.4042916055013 |
| GP2 | Unchanged | 23.4121395072021 | 1.65566876534589 | 25.067808272548 |
| PPY | Increased | 19.917087010905 | 2.85515578139109 | 22.772242792296 |
| SLC30A8 | Increased | 16.3648272475993 | 4.1941028386544 | 20.5589300862537 |
| IAPP | Increased | 15.6884746666667 | 3.08842518721881 | 18.7768998538855 |
| G6PC2 | Unchanged | 17.0355945548096 | 1.20563639040995 | 18.2412309452195 |
| PDX1 | Unchanged | 16.0719715079346 | 1.52676869702853 | 17.5987402049631 |
| UCN3 | Increased | 14.942074 | 2.53161581398156 | 17.4736898139816 |
| SST | Increased | 14.4371310219727 | 2.3272616072825 | 16.7643926292552 |
| ARX | Unchanged | 14.9445376666667 | 1.34593379391941 | 16.2904714605861 |
| NKX2-2 | Unchanged | 15.09123 | 1.16831080102516 | 16.2595408010252 |
| HNF4A | Unchanged | 14.5434013327637 | 1.55499881179239 | 16.0984001445561 |
| ABCC8 | Unchanged | 14.7096996957601 | 1.3383149404851 | 16.0480146362452 |
| FOXA2 | Unchanged | 14.2994064541829 | 1.535513459734 | 15.8349199139169 |
| NEUROD1 | Increased | 12.2341713333333 | 3.54088508710588 | 15.7750564204392 |
| NKX6-1 | Unchanged | 14.1136665445557 | 1.26564618309954 | 15.3793127276552 |
| CHGB | Increased | 12.862944 | 2.44368905706498 | 15.306633057065 |
| PCSK2 | Increased | 12.2508286666667 | 2.94778415781366 | 15.1986128244803 |
| KCNK3 | Unchanged | 12.950375 | 1.14997880619333 | 14.1003538061933 |
| POU5F1 | Unchanged | -10.8860719039714 | 1.91592865247724 | 12.8020005564486 |
| TSPAN1 | Unchanged | 10.8492701692301 | 1.91158085363492 | 12.7608510228651 |
| CHGA | Unchanged | 10.924971597168 | 1.68354297078526 | 12.6085145679532 |
| PCSK1 | Increased | 9.70615746248373 | 2.90216620871523 | 12.608323671199 |
| ISL1 | Unchanged | 10.4585303333333 | 1.47653297459979 | 11.9350633079331 |
| SYP | Unchanged | 10.250111973877 | 1.4618272778411 | 11.711939251718 |
| GLP1R | Increased | 8.76783433333333 | 2.72426794651245 | 11.4921022798458 |
| GCK | Unchanged | 10.2222620024007 | 1.26794205061315 | 11.4902040530139 |
| PAX6 | Increased | 8.7736307006429 | 2.34601684092611 | 11.119647541569 |
| CPE | Increased | 7.58841213155111 | 2.14591785535612 | 9.73432998690722 |
| FGF10 | Increased | 5.484704 | 3.28589663552346 | 8.77060063552346 |
| ITGA1 | Increased | 4.802747 | 3.90286180200824 | 8.70560880200824 |
| MAFA | Unchanged | 7.00004354252116 | 1.66786160118788 | 8.66790514370904 |
| CD274 | Unchanged | 6.6549210333252 | 1.55908587820077 | 8.21400691152597 |
| SOX17 | Unchanged | 6.5393616710612 | 1.60110811237112 | 8.14046978343232 |
| MAFB | Unchanged | 6.42704411344401 | 1.38451189388636 | 7.81155600733037 |
| SLC16A1 | Decreased | -5.21047026985677 | 2.10212971883312 | 7.31259998868989 |
| ONECUT1 | Unchanged | 4.37215382181803 | 1.95343823155986 | 6.32559205337789 |
| IRX2 | Unchanged | 3.86704399808757 | 1.00272368477779 | 4.86976768286536 |
| CXCR4 | Unchanged | 3.12410433333333 | 1.50755299510795 | 4.63165732844127 |
| PAX4 | Unchanged | 2.44286880102539 | 1.87473712275243 | 4.31760592377782 |
| SLC18A1 | Unchanged | 2.44286880102539 | 1.87473712275243 | 4.31760592377782 |
| SOX9 | Unchanged | 3.487971 | 0.58973898079036 | 4.07770998079903 |
| NEUROG3 | Unchanged | -2.81883319897461 | 1.06360164877805 | 3.88243484775266 |
| SLC2A4 | Unchanged | -1.37619183870443 | 0.931327707805937 | 2.30751954651036 |

| Name | Change | Fold change (log2) | Significance | Manhattan distance |
|---|---|---|---|---|
| KCNK1 | Unchanged | 0.487808666666669 | 0.365433355077893 | 0.853242021744562 |
| KRT19 | Unchanged | 0.267876666666666 | 0.426896717364196 | 0.694773384030862 |

### Gene Expression: Volcano Plots Statistics

#### Human Induced Pluripotent Stem Cells versus SC-islets (S6)

### VolcaNoseR - Exploring volcano plots

| Name | Change | Fold change (log2) | Significance | Manhattan distance |
|---|---|---|---|---|
| INS | Increased | 28.646806 | 1.31564790110225 | 29.9624539011023 |
| GCG | Unchanged | 24.5445786666667 | 0.949395858695871 | 25.4939745253625 |
| NEUROD1 | Unchanged | 17.9303113333333 | 0.992041060560456 | 18.9223523938938 |
| PAX4 | Increased | 15.483022 | 1.5392008450508 | 17.0222228450508 |
| TSPAN1 | Unchanged | 15.6127376666667 | 1.10195633595867 | 16.7146940026253 |
| SST | Unchanged | 15.500787 | 0.822953999779912 | 16.3237409997799 |
| SLC18A1 | Increased | 13.8084083518066 | 2.02708094505922 | 15.8354892968659 |
| SLC30A8 | Increased | 14.1690136666667 | 1.5583864523412 | 15.7274001190079 |
| CHGA | Increased | 13.802424 | 1.35088481661801 | 15.153308816618 |
| PDX1 | Unchanged | 13.798629 | 1.13345784233378 | 14.9320868423338 |
| UCN3 | Unchanged | 13.205436 | 1.11220962283364 | 14.3176456228336 |
| ARX | Increased | 12.9127506666667 | 1.33139693239692 | 14.2441475990636 |
| PPY | Unchanged | 12.9382046666667 | 0.829570309406261 | 13.7677749760729 |
| NKX2-2 | Unchanged | 12.091212 | 0.906956895880103 | 12.9981688958801 |
| ISL1 | Unchanged | 11.7262343333333 | 0.899497066000524 | 12.6257313993339 |
| HNF4A | Unchanged | 11.679834 | 0.812492630751022 | 12.492326630751 |
| CHGB | Unchanged | 11.164353 | 1.29708308005213 | 12.4614360800521 |
| NKX6-1 | Increased | 10.922229 | 1.46090372598951 | 12.3831327259895 |
| ABCC8 | Unchanged | 10.9366433333333 | 1.08847449096256 | 12.0251178242959 |
| POU5F1 | Decreased | -10.0401896666667 | 1.85663289885149 | 11.8968225655182 |
| GP2 | Unchanged | 9.83448371496582 | 1.15236318708475 | 10.9868469020506 |
| PAX6 | Unchanged | 9.47139433333333 | 1.09977434733276 | 10.5711686806661 |
| FOXA2 | Unchanged | 9.74264733333333 | 0.79658173157078 | 10.5392290649041 |
| PCSK2 | Increased | 9.07937866666667 | 1.37379725157785 | 10.4531759182445 |
| G6PC2 | Unchanged | 9.324579 | 0.834611935188352 | 10.1591909351884 |
| NEUROG3 | Unchanged | 9.333034 | 0.817905984660298 | 10.1509399846603 |
| MAFB | Unchanged | 8.60953366666666 | 1.11554326687221 | 9.72507693353887 |
| IAPP | Unchanged | 8.66882966666667 | 0.979783941261335 | 9.648613607928 |
| CPE | Unchanged | 8.40225833333333 | 1.10847526073955 | 9.51073359407288 |
| KCNK3 | Unchanged | 7.787368 | 0.914057819862533 | 8.70142581986253 |
| PCSK1 | Increased | 5.86434532869466 | 2.77399845195703 | 8.63834378065169 |
| SOX17 | Unchanged | 7.22220666666667 | 0.742875443400109 | 7.96508211006678 |
| SYP | Unchanged | 6.155397 | 0.943160256153526 | 7.09855725615352 |
| ITGA1 | Increased | 5.705853 | 1.35047790140371 | 7.05633090140371 |
| GLP1R | Unchanged | 6.04482614986165 | 0.916631878010954 | 6.96145802787261 |
| GCK | Unchanged | 5.32745433333334 | 0.828297592038959 | 6.15575192537229 |
| FGF10 | Increased | 4.17222609814453 | 1.49617923812781 | 5.66840533627234 |
| ONECUT1 | Unchanged | 4.74009366666666 | 0.799135251792162 | 5.53922891845883 |
| CXCR4 | Unchanged | 4.52330933333333 | 0.955188268534242 | 5.47849760186757 |
| CD274 | Unchanged | 3.93775604003906 | 1.00755356133271 | 4.94530960137177 |
| KCNK1 | Unchanged | 3.63340466666666 | 1.02238205822919 | 4.65578672489586 |
| MAFA | Unchanged | 3.24972066666666 | 0.999740859942211 | 4.24946152660888 |
| KRT19 | Unchanged | 2.20293766666666 | 0.908884402230925 | 3.11182206889759 |
| SOX9 | Unchanged | -2.201004 | 0.879089361508815 | 3.08009336150882 |
| SLC16A1 | Unchanged | -1.34429747932943 | 1.50078651201772 | 2.84508399134715 |

| Name | Change | Fold change (log2) | Significance | Manhattan distance |
|---|---|---|---|---|
| IRX2 | Unchanged | 1.71674301847331 | 0.339327357201652 | 2.05607037567496 |
| SLC2A4 | Unchanged | 0.240448252237956 | 0.777084398287654 | 1.01753265052561 |

Gene Expression: Volcano Plots Statistics

SC-islets (S6) versus Grafts

### VolcaNoseR - Exploring volcano plots

| Name | Change | Fold change (log2) | Significance | Manhattan distance |
|---|---|---|---|---|
| GCG | Unchanged | 24.1648916075099 | 0.840971862421787 | 25.0058634699317 |
| NEUROD1 | Unchanged | 17.9022125207357 | 0.98126117837118 | 18.8834736991069 |
| PAX4 | Increased | 15.4828507135301 | 1.53907133695608 | 17.0219220504862 |
| TSPAN1 | Unchanged | 15.5586188656051 | 1.06258111665318 | 16.6211999822583 |
| SLC18A1 | Increased | 13.8078614762308 | 2.02656826013198 | 15.8344297363628 |
| SST | Unchanged | 14.5617525781634 | 0.714140342909831 | 15.2758929210733 |
| CHGA | Unchanged | 13.5913903763786 | 1.18662189272567 | 14.7780122691043 |
| POU5F1 | Unchanged | -11.2127623709995 | 0.728881189932064 | 11.9416435609316 |
| ISL1 | Unchanged | 10.9519529496917 | 0.742218910993392 | 11.6941718606851 |
| NEUROG3 | Unchanged | 9.33271693687875 | 0.818822571570704 | 10.1515395084495 |
| MAFB | Unchanged | 8.25053345189797 | 0.964121155314509 | 9.21465460721248 |
| PAX6 | Unchanged | 8.0885906214998 | 0.592907629919154 | 8.68149825141896 |
| CPE | Unchanged | 7.18847576695779 | 0.611433751512922 | 7.79990951847071 |
| SOX17 | Unchanged | 5.81508939269419 | 0.727872006189134 | 6.54296139888332 |
| ITGA1 | Unchanged | 4.60197988247833 | 0.696314670471084 | 5.29829455294941 |
| CXCR4 | Unchanged | 3.83565385183593 | 0.824150982801238 | 4.65980483463717 |
| KCNK1 | Unchanged | 3.460409488687 | 1.01360712664215 | 4.47401661532915 |
| ONECUT1 | Unchanged | 2.58880699594552 | 0.725982366415262 | 3.31478936236078 |
| SLC30A8 | Unchanged | 0 | 3.27617482636514 | 3.27617482636514 |
| SLC16A1 | Decreased | -1.44678601897163 | 1.80167618206771 | 3.24846220103934 |
| IAPP | Unchanged | 0 | 3.06448126098871 | 3.06448126098871 |
| PPY | Unchanged | 0 | 2.7977829631042 | 2.7977829631042 |
| PCSK1 | Unchanged | 0 | 2.77177893615941 | 2.77177893615941 |
| KRT19 | Unchanged | 1.76558761030835 | 0.985533966663901 | 2.75112157697225 |
| PCSK2 | Unchanged | 0 | 2.4766100749648 | 2.4766100749648 |
| FGF10 | Unchanged | 0 | 1.71728079438097 | 1.71728079438097 |
| GP2 | Unchanged | 0 | 1.65550836036037 | 1.65550836036037 |
| MAFA | Unchanged | 0 | 1.525432691174 | 1.525432691174 |
| CD274 | Unchanged | 0 | 1.34594611669173 | 1.34594611669173 |
| CHGB | Unchanged | 0 | 1.34413637399991 | 1.34413637399991 |
| GLP1R | Unchanged | 0 | 1.33983410874936 | 1.33983410874936 |
| SYP | Unchanged | 0 | 1.27777879559632 | 1.27777879559632 |
| SLC2A4 | Unchanged | -0.32893252408168 | 0.915795673900126 | 1.24472819798181 |
| G6PC2 | Unchanged | 0 | 1.19096059997675 | 1.19096059997675 |
| ABCC8 | Unchanged | 0 | 1.18867871366603 | 1.18867871366603 |
| PDX1 | Unchanged | 0 | 1.1801552705372 | 1.1801552705372 |
| FOXA2 | Unchanged | 0 | 1.16872578532968 | 1.16872578532968 |
| GCK | Unchanged | 0 | 1.15152640411998 | 1.15152640411998 |
| NKX6-1 | Unchanged | 0 | 1.12686372504837 | 1.12686372504837 |
| KCNK3 | Unchanged | 0 | 1.10333213393286 | 1.10333213393286 |
| SOX9 | Unchanged | 0 | 1.08056222910341 | 1.08056222910341 |
| IRX2 | Unchanged | 0 | 1.02967593077903 | 1.02967593077903 |
| NKX2-2 | Unchanged | 0 | 0.879983207674042 | 0.879983207674042 |
| UCN3 | Unchanged | 0 | 0.794824274326348 | 0.794824274326348 |
| HNF4A | Unchanged | 0 | 0.718821223726417 | 0.718821223726417 |

| Name | Change | Fold change (log2) | Significance | Manhattan distance |
|---|---|---|---|---|
| ARX | Unchanged | 0 | 0.653648384452424 | 0.653648384452424 |
| INS | Unchanged | 0 | 0.583906844858268 | 0.583906844858268 |
| GAPDH | Unchanged | 0 | 0 | 0 |

Gene Expression: Volcano Plots Statistics

SC-islets (S6) versus Human Donor Cadaveric Islets

### VolcaNoseR - Exploring volcano plots

| Name | Change | Fold change (log2) | Significance | Manhattan distance |
|---|---|---|---|---|
| PCSK1 | Increased | 2.90712398765089 | 6.14941596234189 | 9.05653994999278 |
| INS | Increased | 3.65052802315504 | 4.09032411413389 | 7.74085213728893 |
| CHGA | Increased | 3.39399640551225 | 3.79205639609054 | 7.18605280160279 |
| SLC30A8 | Increased | 2.82500219631564 | 3.60898563481198 | 6.43398783112762 |
| ARX | Increased | 3.58990601683874 | 2.73961459753479 | 6.32952061437354 |
| ABCC8 | Increased | 3.22981153637247 | 3.0077376772823 | 6.23754921365477 |
| SLC18A1 | Increased | 3.46743474430475 | 2.29630844635201 | 5.76374319065675 |
| PDX1 | Increased | 2.97537107999145 | 2.45020623308924 | 5.42557731308069 |
| NKX2-2 | Unchanged | 3.86483398635174 | 1.52776763966676 | 5.3926016260185 |
| CHGB | Increased | 2.15803435155458 | 3.1970700833425 | 5.35510443489708 |
| PAX4 | Increased | 3.07946335954374 | 2.25976651491642 | 5.33922987446016 |
| ISL1 | Increased | 2.36216123436111 | 2.83148787258964 | 5.19364910695075 |
| GLP1R | Increased | 2.77792523688345 | 2.16566709792105 | 4.9435923348045 |
| FOXA2 | Unchanged | 3.12733076914808 | 1.79892191356479 | 4.92625268271287 |
| ITGA1 | Increased | 1.97194702492214 | 2.35773867558581 | 4.32968570050795 |
| GCK | Unchanged | 3.0838511435183 | 1.21698741668085 | 4.30083856019915 |
| MAFB | Increased | 1.55644066084296 | 2.61889257896213 | 4.17533323980509 |
| GCG | Increased | 1.93781713652475 | 2.12128213924652 | 4.05909927577127 |
| NEUROD1 | Increased | 1.5984791051547 | 2.26112654892819 | 3.85960565408288 |
| HNF4A | Unchanged | 3.02861290232584 | 0.715966400278465 | 3.74457930260431 |
| G6PC2 | Unchanged | 2.2594540669206 | 1.40785208470301 | 3.66730615162361 |
| PCSK2 | Unchanged | 1.52859285754013 | 1.97917930305782 | 3.50777216059796 |
| CPE | Unchanged | -1.98881847640665 | 1.45732058482546 | 3.44613906123211 |
| ONECUT1 | Unchanged | 1.39336461457519 | 1.83573282954949 | 3.22909744412468 |
| NKX6-1 | Unchanged | 1.82446514364972 | 1.34440338637785 | 3.16886853002758 |
| IRX2 | Unchanged | 1.69711234305846 | 1.45730508779229 | 3.15441743085075 |
| SYP | Unchanged | 1.95515286064845 | 0.91952783881041 | 2.87468069945886 |
| KCNK3 | Unchanged | 2.55133472327131 | 0.32252467920878 | 2.87385940248009 |
| IAPP | Unchanged | 0.738832902956974 | 1.49538131521517 | 2.23421421817214 |
| PAX6 | Unchanged | -0.391279271073528 | 1.51930277725095 | 1.91058204832448 |
| CD274 | Unchanged | 1.30551134051293 | 0.384426439166712 | 1.68993777967965 |
| PPY | Unchanged | -0.64200270063364 | 0.917522840930397 | 1.55952554156404 |
| FGF10 | Unchanged | -1.37770686616395 | 0.0173584410314411 | 1.3950653071954 |
| SST | Unchanged | 0 | 1.24905883694906 | 1.24905883694906 |
| CXCR4 | Unchanged | 0 | 1.20355388354402 | 1.20355388354402 |
| GP2 | Unchanged | 0 | 1.00163562109317 | 1.00163562109317 |
| UCN3 | Unchanged | 0 | 0.976472895929325 | 0.976472895929325 |
| SOX17 | Unchanged | 0 | 0.807433805563026 | 0.807433805563026 |
| KRT19 | Unchanged | 0.734349236145854 | 0.00596040052489852 | 0.740309636670753 |
| TSPAN1 | Unchanged | 0 | 0.524137207846251 | 0.524137207846251 |
| SOX9 | Unchanged | 0 | 0.317297457559239 | 0.317297457559239 |
| MAFA | Unchanged | 0 | 0.227404829732896 | 0.227404829732896 |
| KCNK1 | Unchanged | 0 | 0.18879070213504З | 0.188790702135043 |
| NEUROG3 | Unchanged | 0 | 0.145278235410755 | 0.145278235410755 |
| GAPDH | Unchanged | 0 | 0 | 0 |

| Name | Change | Fold change (log2) | Significance | Manhattan distance |
|---|---|---|---|---|
| SLC16A1 | Unchanged | 0 | 0 | 0 |
| SLC2A4 | Unchanged | 0 | 0 | 0 |
| POU5F1 | Unchanged | 0 | 0 | 0 |
