## Supplementary material for "Complete Suspension Differentiation of Human Pluripotent Stem Cells into Pancreatic Islets Using Vertical Wheel^®^ Bioreactors": Tables S1-7

**Table S1. Patient demographics used in this study.**

| iPSC line | Age | Sex | Gender | Health status |
|---|---|---|---|---|
| **#1** | 53 | Female | Female | Healthy |
| **#2** | 43 | Female | Female | Diabetic |
| **#3** | 30 | Male | Male | Diabetic |

**Table S2. Quality control parameters for screening induced pluripotent stem cells.**

| Attribute | Test | Analytical method | Acceptance criteria |
|---|---|---|---|
| Microbiological sterility | Mycoplasma | qPCR | Negative |
| Genetic fidelity and stability | Viral testing | PCR | Negative |
| | Karyotype | qPCR | Copy number = 2 |
| Characterization | Immunocytochemistry | A minimum of two pluripotency markers (Sox2, Oct4) | Both markers should be positive on > 90% of cells. |
| | Flow cytometry | A minimum of three pluripotency markers (Sox2, Oct4, Nanog). | All markers should be positive on > 90% of cells. |
| Potency | Phenotypic | Teratoma formation | Demonstration of cells from all three germ layers. |

*Table S3. Differentiation medium composition.*

| Stage | Duration (d) | Basal medium | Additives |
|---|---|---|---|
| **1** | 4 | STEMdiff Definitive Endoderm Basal Medium (STEMCell Technologies, cat. 05111) | 1% (v/v) CJ (STEMCell Technologies, cat. 05113)<br>1% (v/v) MR (STEMCell Technologies, cat. 05112) on the first day. |
| **2** | 2 | RPMI 1640 (Gibco, cat. 61870-036)<br>1% (v/v) Glutamax (Gibco, cat. A12860-01) | 1% (v/v) B-27 Serum-free supplement (50x) (Life Technology, cat. 17504044)<br>50 ng/ml FGF7 (R&D Systems, cat. 251-KG) |
| **3** | 2 | DMEM (Gibco, cat. 10569-010)<br>1% (v/v) Glutamax<br>1mM Sodyum Pyruvate (Gibco, cat. 11360070)<br>10 mM HEPES (Gibco, cat. 15630080) | 1% (v/v) B-27 Serum-free supplement (50x)<br>0.25 µM KAAD-Cyclopamine (EMD Millipore, cat. 239804)<br>2 µM Retinoic acid (STEMCell Technologies, cat. 73794)<br>0.25 µM LDN193189 (Stemgent, cat. 04-0074) |
| **4** | 4 | DMEM<br>1% (v/v) Glutamax<br>1mM Sodyum Pyruvate<br>10 mM HEPES | 1% (v/v) B-27 Serum-free supplement (50x)<br>25 ng/ml FGF7<br>50 ng/ml EGF (R&D Systems, cat 236-EG) |
| | 1 | | 1% (v/v) B-27 Serum-free supplement (50x)<br>25 ng/ml FGF7<br>10 µg/ml Heparin (Sigma, cat. H3149)<br>10 µM RockI (STEMCell Technologies, cat. 72304)<br>1 µM Aphidicolin (Sigma, cat. 5.04744.0001) |
| **5** | 7 | RPMI 1640<br>1% (v/v) Glutamax | 1% (v/v) B-27 Serum-free supplement (50x)<br>1 µM thyroid hormone (T3) (Cat. No. T6397, Sigma)<br>10 µM ZnSO4 (Sigma, cat. Z0251-100G)<br>10 µg/ml Heparin<br>10 µM RockI<br>10 µM ALK5 inhibitor<br>1 µM Aphidicolin (Sigma, cat. 5.04744.0001) |
| **6** | 7 | RPMI 1640<br>1% (v/v) Glutamax | 1% (v/v) B-27 Serum-free supplement (50x)<br>10% (v/v) KnockOut serum replacement (Life Technologies, cat. 10828-028)<br>1 µM Thyroid hormone (T3)<br>10 µM ZnSO4<br>10 µg/ml Heparin<br>10 µM RockI |

**Table S4. Patient demographic for human islet donors**

| Donor ID | Age | Sex | Purity | BMI | Diabetes | HbA1c (%) | IEQ |
|---|---|---|---|---|---|---|---|
| R408 | 39 | M | 80% | 27.5 | Non-diabetic | 5.6 | 10,000 |
| R410 | 59 | F | 80% | 26.4 | Non-diabetic | 4.8 | 12,000 |
| R427 | 52 | M | 90% | 35.8 | Non-diabetic | 5.8 | 25,000 |

**Table S5. Quantitative Polymerase Chain Reaction Sequence for Karyotype Analysis**

| **Stage** | **Cycles** | **Temperature (°C)** | **Cycling Time (min:sec)** |
|---|---|---|---|
| **Polymerase Activation** | 1 | 95.0 | 3:00 |
| **Denature** | 40 | 95.0 | 0:05 |
| **Anneal** | | 60.0 | 0:30 |

**Table S6. Antibodies and concentrations used for immunohistochemistry (IHC) and flow cytometry (FC).**

*All secondaries for immunohistochemistry were applied at a 1:250 concentration and all secondaries for flow cytometry were applied at a 1:500 concentration.

| **Epitope** | **Origin animal** | **Conjugate** | **Dilution** | **Supplier** | **Assay** |
|---|---|---|---|---|---|
| Oct4 | Mouse | FITC | 1:100 | Sigma (cat. MAB4419A4) | IHC |
| Oct4 | Mouse | BV421 | 1:100 | BD (cat. 565644) | FC |
| Sox2 | Mouse | PE | 1:100 | Biorbyt (cat. orb124865) | IHC |
| CD184 | Mouse | BV421 | 1:100 | BD (cat. 562448( | FC |
| CD117 | Mouse | FITC | 1:50 | Invitrogen (cat. 11-1178-42) | FC/ IHC |
| SOX17 | Mouse | APC | 1:20 | R&D Systems (cat. IC1924A) | FC/ IHC |
| FoxA2 | Rabbit | N/A | 1:100 | Abcam (cat. 108422) | FC/ IHC |
| Pdx1 | Mouse | FITC | 1:20 | BD (cat. 562274) | FC |
| Pdx1 | Goat | N/A | 1:20 | R&D Systems (cat. AF2419) | IHC |
| Nkx6.1 | Mouse | N/A | 1:10 | DSHB (cat. F55A10-c) | FC/ IHC |
| Chromogranin A | Mouse | AF405 | 1:50 | Novus (NBP2-33198AF405) | FC/ IHC |
| GP2 | Mouse | AF405 | 1:50 | Novus (cat. NBP3-08243AF405) | FC/ IHC |
| C-peptide | Rabbit | N/A | 1:100 | Abcam (cat. ab14181) | FC/ IHC |
| Isl1 | Mouse | PE | 1:50 | BD (cat. Q11-465) | IHC |
| Insulin | Guinea Pig | N/A | 1:500 | DAKO (cat. A0564) | IHC |

| | | | | | |
|---|---|---|---|---|---|
| Glucagon | Mouse | N/A | 1:800 | Sigma (cat. G2654) | IHC |
| UCN3 | Rabbit | N/A | 1:100 | Abbexa (cat. abx100886) | FC/ IHC |
| Somatostatin | Mouse | PE | 1:100 | R&D Systems (cat. FAB4224P) | IHC |
| Grehlin | Rabbit | N/A | 1:50 | BioVision (cat. 5991-100) | IHC |
| Ki67 | Rabbit | N/A | 1:50 | Abcam (cat. ab15580) | IHC |
| SLC18A1 | Rabbit | N/A | 1:100 | Sigma (cat. HPA063797) | FC |
| Nkx2.2 | Mouse | N/A | 1:20 | DSHB (cat. 74.5A5) | IHC |
| Pancreatic polypeptide | Rabbit | N/A | 1:50 | Abcam (cat. Ab14985) | IHC |
| Ck19 | Rabbit | N/A | 1:50 | Abcam (cat. ab52625) | IHC |
| Anti-goat | Donkey | FITC | - | Thermo Fisher (cat. A16000) | FC/ IHC |
| Anti-mouse | Donkey | AF647 | - | Invitrogen (cat. A31571) | FC/ IHC |
| Anti-mouse | Goat | PE | - | Jackson (cat. 115-115-164) | FC/ IHC |
| Anti-rabbit | Donkey | AF405 | - | Invitrogen (cat. A48258) | FC/ IHC |
| Anti-rabbit | Goat | AF594 | - | Invitrogen (cat. A11012) | FC/ IHC |
| Anti-guinea pig | Goat | FITC | - | Invitrogen (cat. A11073) | FC/ IHC |

### Table S7. Thermo Fisher TaqMan Micro Fluidic Array Card Configuration

| Assay ID | Gene | Gene Name(s) | Amplicon Length |
|---|---|---|---|
| Hs01093752_m1 | ABCC8 | ATP binding cassette subfamily C member 8 | 58 |
| Hs00292465_m1 | ARX | Aristaless related homeobox | 96 |
| Hs00900370_m1 | CHGA | Chromogranin A | 67 |
| Hs01084631_m1 | CHGB | Chromogranin B | 112 |
| Hs00175676_m1 | CPE | Carboxypeptidase E | 106 |
| Hs00607978_s1 | CXCR4 | C-X-C motif chemokine receptor 4 | 153 |
| Hs00204257_m1 | CD274 | CD274 molecule | 77 |
| Hs00610298_m1 | FGF10 | Fibroblast growth factor 10 | 70 |
| Hs00232764_m1 | FOXA2 | Forkhead box A2 | 66 |
| Hs01549772_m1 | G6PC2 | Glucose-6-phosphatase catalytic subunit 2 | 97 |
| Hs99999905_m1 | GAPDH | Glyceraldehyde-3-phosphate dehydrogenase | 0 |

| | | | |
|---|---|---|---|
| Hs01031536_m1 | GCG | Glucagon | 86 |
| Hs01564555_m1 | GCK | Glucokinase | 72 |
| Hs00157705_m1 | GLP1R | Glucagon like peptide 1 receptor | 78 |
| Hs00230853_m1 | HNF4A | Hepatocyte nuclear factor 4 alpha | 49 |
| Hs00846499_s1 | UCN3 | Urocortin 3 | 85 |
| Hs00355773_m1 | INS | Insulin | 126 |
| Hs01383002_m1 | IRX2 | Iroquois homeobox 2 | 85 |
| Hs00158126_m1 | ISL1 | ISL LIM homeobox 1 | 57 |
| Hs00235006_m1 | ITGA1 | Integrin subunit alpha 1 | 87 |
| Hs01116799_m1 | KCNK1 | Potassium two pore domain channel subfamily K member 1 | 140 |
| Hs00605529_m1 | KCNK3 | Potassium two pore domain channel subfamily K member 3 | 134 |
| Hs00761767_s1 | KRT19 | Keratin 19 | 116 |
| Hs04419852_s1 | MAFA | MAF bzip transcription factor A | 107 |
| Hs00534343_s1 | MAFB | MAF bzip transcription factor B | 86 |
| Hs01922995_s1 | NEUROD1 | Neuronal differentiation 1 | 110 |
| Hs01875204_s1 | NEUROG3 | Neurogenin 3 | 127 |
| Hs00159616_m1 | NKX2-2 | NK2 homeobox 2 | 114 |
| Hs00232355_m1 | NKX6-1 | NK6 homeobox 1 | 93 |
| Hs00413554_m1 | ONECUT1 | One cut homeobox 1 | 76 |
| Hs00173014_m1 | PAX4 | Paired box 4 | 115 |
| Hs00240871_m1 | PAX6 | Paired box 6 | 76 |
| Hs01026107_m1 | PCSK1 | Proprotein convertase subtilisin/kexin type 1 | 96 |
| Hs00159922_m1 | PCSK2 | Proprotein convertase subtilisin/kexin type 2 | 76 |
| Hs00236830_m1 | PDX1 | Pancreatic and duodenal homeobox 1 | 73 |
| Hs00426805_m1 | GP2 | Glycoprotein 2 | 75 |
| Hs00358111_g1 | PPY | Pancreatic polypeptide | 68 |
| Hs01560299_m1 | SLC16A1 | Solute carrier family 16 member 1 | 95 |

| | | | |
|---|---|---|---|
| Hs00915193_m1 | SLC18A1 | Solute carrier family 18 member A1 | 63 |
| Hs00168966_m1 | SLC2A4 | Solute carrier family 2 member 4 | 89 |
| Hs00545183_m1 | SLC30A8 | Solute carrier family 30 member 8 | 73 |
| Hs00751752_s1 | SOX17 | SRY-box 17 | 149 |
| Hs00165814_m1 | SOX9 | SRY-box 9 | 102 |
| Hs00356144_m1 | SST | Somatostatin | 86 |
| Hs00300531_m1 | SYP | Synaptophysin | 63 |
| Hs00169095_m1 | IAPP | Islet amyloid polypeptide | 61 |
| Hs00371661_m1 | TSPAN1 | Tetraspanin 1 | 87 |
| Hs04260367_gH | POU5F1 | POU class 5 homeobox 1 | 77 |
